# Screening of anti-*Acinetobacter baumannii* phytochemicals, based on the potential inhibitory effect on OmpA and OmpW functions

**DOI:** 10.1101/2021.07.10.451884

**Authors:** Shahab Shahryari, Parvin Mohammadnejad, Kambiz Akbari Noghabi

## Abstract

Therapeutic options, including last-line or combined antibiotic therapies for multi-drug resistant (MDR) strains of *Acinetobacter baumannii* are ineffective. The outer membrane protein A (OmpA) and outer membrane protein W (OmpW) are two porins known for their different cellular functions. Identification of natural compounds with the potentials to block these putative porins can attenuate the growth of the bacteria and control the relating diseases. The current work aimed to screen a library of 384 phytochemicals according to their potentials to be used as a drug, and potentials to inhibit the function of OmpA and OmpW in *A. baumannii*. The phytocompounds were initially screened based on their physicochemical, absorption, distribution, metabolism, excretion, and toxicity (ADMET) drug-like properties. Afterward, the selected ligands were subjected to standard docking calculations against the predicted three-dimensional structure of OmpA and OmpW in *A. baumannii*. We identified three phytochemicals (isosakuranetin, aloe-emodin and pinocembrin) possessing appreciable binding affinity towards the selected binding pocket of OmpA and OmpW. Molecular dynamics (MD) simulation analysis confirmed the stability of the complexes. Amongst them, isosakuranetin was suggested as the best phytocompound for further *in vitro* and *in vivo* study.

## 1. Introduction

In recent years, antimicrobial resistance (AMR) by Gram-negative bacteria has become a serious problem with a possible devastating impact on the economy and human life [1]. With the advent of the post-antibiotic era, the need to use more effective and safer antimicrobial compounds is fully felt. [2]. As one of the six superbug ESKAPE pathogens, *Acinetobacter baumannii* is a naturally transformable Gram-negative pathogen with increasing clinical importance [3]. Multi-drug resistant (MDR) strains of *Acinetobacter* are responsible for chronic infections, and can rapidly confer resistance to the most of currently used antibiotics. Thus, *A. baumannii* is in the global priority pathogen list (PPL) to develop effective antimicrobial therapies against it [4]. Due to the rapid changes in the genetic constitution of *A. baumannii* and lack of enough knowledge about resistance mechanisms, no efficient antibiotic against this microorganism has yet been developed by the pharmaceutical industry [5]. This paucity of effective antibiotics has revived scientific interest in finding efficient antimicrobial agents, with the ability to kill, inhibit the growth, or inhibit the activity of some essential virulence factors of *A. baumannii* [6-9].

The outer membrane proteins (OMPs) of Gram-negative bacteria, have attracted some attention as drug targets due to their extensive functions [10, 11]. Outer membrane protein A (OmpA) as the most abundant slow porin in members of genus *Acinetobacter* plays different rules in cells [12, 13]. This protein is closely associated with the virulence and survival of the bacteria under harsh conditions [14-16]. OmpA structure is composed of a β-barrel and a periplasmic (OmpA-like) domain. In addition to a structural role for OmpA of *A. baumannii*, it can play an important role in biofilm formation and adhesion to host cells [12, 17-21]. The absence of OmpA in *A. baumannii* and *E. coli* is associated with significant reduction of virulence [22, 23]. Also, it has been suggested that 4 external loops, connecting N-terminal β-strands, participate in some interactions with host-cells (or other surfaces), and can be used as drug-targets [24].

Outer membrane protein W (OmpW) is another porin with a pivotal role in the uptake of nutritional substances such as iron [25]. The role of iron acquisition in the pathogenicity of *A. baumannii* has been reported previously [26], and disruption of this system can be effective in treating infections caused by this microorganism [27-29]. Decreased minimum inhibitory concentrations (MICs) of several antibiotics due to deletion of OmpW from *E. coli* [30], as well as reduced intestinal colonization of *ΔompW* mutant strains of *V. cholerae* [31], are among the indications that OmpW can be served as a potential drug target in Gram-negative bacteria. *In silico* and *in vitro* studies on OmpW resulted in the identification of three inhibitors of this porin in *A. baumannii* [32]. Therefore, any molecule with the ability to disrupt the functions of OmpA and OmpW can help treat the disease caused by this bacterium.

Researchers have reported the benefits of infectious disease treatment using poly-pharmacological drugs [33]. In light of this, the aim of the current study was *in silico* screening of the plant based compounds with potential anti-*A. baumannii* activities. The adhesion property of OmpA and the function of OmpW in the uptake of nutrients were targeted for the screening of compounds. For this purpose, a library consisting of 384 plant-based compounds, known as an anti-bacterial or anti-viral compound, was selected for this study [9, 34-36]. We applied *in silico* approaches to prioritize these biomolecules for toxicological evaluation and safety concerns [37, 38]. The phytocompounds with the highest affinity for both OmpA and OmpW were selected as the multi-target drug candidates. The results for molecular docking were further evaluated using molecular dynamics (MD) simulation. Our results led to the identification of three potential mutual OmpA and OmpW functional inhibitors, isosakuranetin, aloe-emodin and pinocembrin. Isosakuranetin was suggested for further *in vitro* and *in vivo* studies.

## 2. Data and methodology

### 2.1. Tools and servers

A list of all tools and servers used in this work with a brief description is illustrated in Table S1.

### 2.2. Targets sequences

The complete amino acid sequences of OmpA (BAN86529 accession number) and OmpW (BAN89067 accession number) were retrieved from the GenBank database at National Center for Biotechnology Information (NCBI) (http://www.ncbi.nlm.nih.gov/protein/) in FASTA format.

### 2.3. Signal peptide prediction

The secretary nature of selected proteins was predicted using SignalP 4.1 server (http://www.cbs.dtu.dk/services/SignalP/), which integrates a prediction of cleavage sites and signal or non-signal peptide prediction based on a combination of several artificial neural networks. LipoP 1.0 server (http://www.cbs.dtu.dk/services/LipoP) used to predict probable signal peptide within the sequence and signal peptidase I or II cleavage sites within the protein.

### 2.4. Homology Modeling, model refinement, and energy minimization

As the full three-dimensional (3D) structure of none of the proteins was available in the Protein Data Bank (PDB), a homology modeling was performed to predict the 3D structure of each protein using RaptorX web server (http://raptorx.uchicago.edu/StructurePrediction/predict/). The server predicts the absolute global quality and comparable global quality for each of the residues of the query sequence. In order to check the 3D structure, we used PyMOL molecular graphics system version 1.7.4.4 (Schrödinger, LLC, and Portland, OR, USA). To perform structure refinement and energy minimization of the 3D modeled protein structures, the online server GalaxyRefine (http://galaxy.seoklab.org/) and YASARA software were used, respectively. GalaxyRefine employs the CASP10 assessment to refine the query structure, improving the structural and global quality of the 3D model. This method initially rebuilds side chains and performs side chain repacking and subsequently uses molecular dynamics simulation to achieve overall structure relaxation.

### 2.5. Validation of the 3D structures

To validate the refined and optimized 3D structures, three freely available web tools of RAMPAGE (http://mordred.bioc.cam.ac.uk/rapper/~rampage.php), ProSA-web (https://prosa.services.came.sbg.ac.at/prosa.php), and ERRAT (https://servicesn.mbi.ucla.edu/ERRAT/) were used. The server RAMPAGE evaluates the Ramachandran plot by applying PROCHECK principles. The ProSA validation method evaluates model accuracy and statistical significance with a knowledge-based potential. It plots overall excellence scores of the errors calculated in the query 3D structure. Further, the ERRAT server was used to check the reliability of modeled proteins and their overall quality factors.

### 2.6. Prediction of potential ligand binding pocket

To avoid blind docking and increase the accuracy of molecular docking, the top three possible binding sites of the proteins were identified using METAPOCKET version 2.0 (https://projects.biotec.tu-dresden.de/metapocket/index.php). As a META server, METAPOCKET is a popular consensus method, which combines eight methods of asLIGSITEcs, PASS, QSiteFinder, SURFNET, Fpocket, GHECOM, Concavity and POCASA to predict binding sites based on 3D protein structures.

### 2.7. Natural-compound ligand library

A library of 384 phytochemicals (Table S2) with potential anti-infection properties was used in this study [9, 34-36, 39]. The 3D structures of the molecules were retrieved from UniProt darabase. In the case of lacking X-ray crystallography or NMR structure of the compound, a 3D conformer was obtained from PubChem database. The ligands 3D structures were energy minimized using Chem3D software before docking. The ligand files were finally saved in PDB format for molecular docking studies.

### 2.8. Pre-filtration and pharmacokinetic analysis of phytochemicals

A compound must pass through multiple filters to be considered as a novel drug. All of the bioactive candidates used in this study were evaluated based on their important physicochemical properties using SwissADME (http://www.swissadme.ch/) online server. Initially, those compounds satisfying Lipinski’s rule of five (RO5) were selected and those that violated the rule were eliminated from downstream analysis. Pharmacokinetic properties of absorption, distribution, metabolism, excretion, and toxicity (ADMET) with a crucial role in the development of drug design were predicted using the pkCSM webserver at (http://biosig.unimelb.edu.au/pkcsm/) to decrease the failure rate of the compound for further analysis. For further screening PreADME (https://preadmet.bmdrc.kr/) server was used to filter the compounds based on their drug-likeliness, ADME properties and toxicity.

### 2.9. Molecular docking analysis

Molecular docking analysis was performed using AutoDock Vina software [40]. The .pdbqt file of ligands was prepared after specifying of rotatable, non-rotatable bonds, and torsion angles in the compounds using AutoDock tools. Grid boxes were defined according to the selected binding pockets and the aim of this study. The protein (receptor) files in .pdbqt format were prepared for a flexible body docking using AutoDock Tools software. As the target for OmpA was external loops because of their potential for adhesion, all the residues of external loops in the predicted binding pocket, were chosen for torsions. Concerning OmpW, only the residues of the binding pocket located toward the β-barrel were selected for torsion, because of the aim of this study for screening of compounds with the ability to block the OmpW channel. The docking process was performed after preparing a configuration file. The obtained interactions and generated poses were visualized using PyMOL. The docking results were evaluated for their orientation, number of hydrogen bonds, number of interacting residues, binding affinity (kcal/mol), and RSMD.

### 2.10. Drug-Target Interactions and Mechanism of Binding

The mechanism of binding in drug-target complexes was profiled using the LigPlot^+^ program (https://www.ebi.ac.uk/thornton-srv/software/LigPlus/). Hydrogen bonds, as well as hydrophobic interactions between potential drug molecules and the protein target were studied via 2D interaction diagrams of the docked drug-target complexes.

### 2.11. BOILED-Egg Model for Prediction of gastrointestinal absorption (GI) and blood-brain barrier penetration (BBB)

To reveal the capability of GI absorption and permeability of the Blood-Brain Barrier (BBB), the BOILED-Egg model of the potential drugs was predicted using SwissADME (http://www.swissadme.ch/).

### 2.12. Molecular dynamics (MD) simulation

MD simulation was used to refine the 4 complexes obtained from molecular docking simulation. MD analysis was performed using a GROMACS 4.6.5 with GROMOS96 force field and simple point charge (SPC) as a water model [41].

To mimic the experimental conditions the simulation was carried out at 343 K. Some molecules of water were randomly replaced by ions for neutralization. All systems were simulated under the NpT ensemble within periodic boundary conditions. The neighbor list was updated with 10 step frequency using the grid search method. The leap-frog algorithm was utilized with a 2 fs time step. Protein and solvent were coupled separately to a heat bath at the desired temperature with time constant τT = 0.1 ps applying V-rescale thermostat [42]. After being neutralized, the system was submitted to energy minimization applying the steepest descent algorithm. After that, the equilibrations of systems were done under NVT up to 100 ps at 300 K with restraint forces of 1000 kJ/mol, followed by 100 ps under NPT at the pressure of 1 bar and with restraint forces of 1000 kJ/mol using modified Berendsen thermostat and Parinello-Rahman barostat algorithms respectively. The electrostatic interactions were analyzed using the Particle Mesh Ewald (PME) method .Ultimately, the MD run was performed with no restraint for 50 ns.

## 3. Results and discussion

### 3.1. Homology modeling, model refinement, and energy minimization

Multiple templates were used to predict the 3D structure of OmpA and OmpW (Figure 1 and 2). Crystal structure of OmpA-like domain from *A. baumannii* (3TD3) with p-value 6.35e-06 was used as a template for prediction of OmpA periplasmic domain (Figure 1). Five β-barrel crystal structures (1P4T, 2X4M, 3NB3, 3QRA, and 4RLC) were suggested for3D structure prediction of this domain in *A. baumannii* OmpA (Figure 1). The crystal structure 4RLC with p-value 2.05e-14 was used as the best template. The tertiary structure of OmpW predicted to contain only one domain)β-barrel domain) and crystal structure 2F1T with p-value 3.48e-07 was employed as the best template for this prediction (Figure 2). Refinement of the obtained “crude” models from the GalaxyRefine server resulted in five models for each protein (Table S3). Based on model quality scores for all refined models, model two with GDT-HA value of 0.9633, RMSD of 0.384, and MolProbity of 1.922, clash score of 17.7, and poor rotamers score of 0.7 was found to be the best model of OmpA. Its Ramachandran plot score was 97.0%. For OmpW, model three was found to be the best model based on GDT-HA of 0.9684, RMSD of 0.392, MolProbity of 1.746, clash score of 11.8, and poor rotamers of 0.0. Ramachandran plot score was 97.1% for this model. YASARA energy minimization tool was used to further improve the quality of the models.

**Figure 1.**
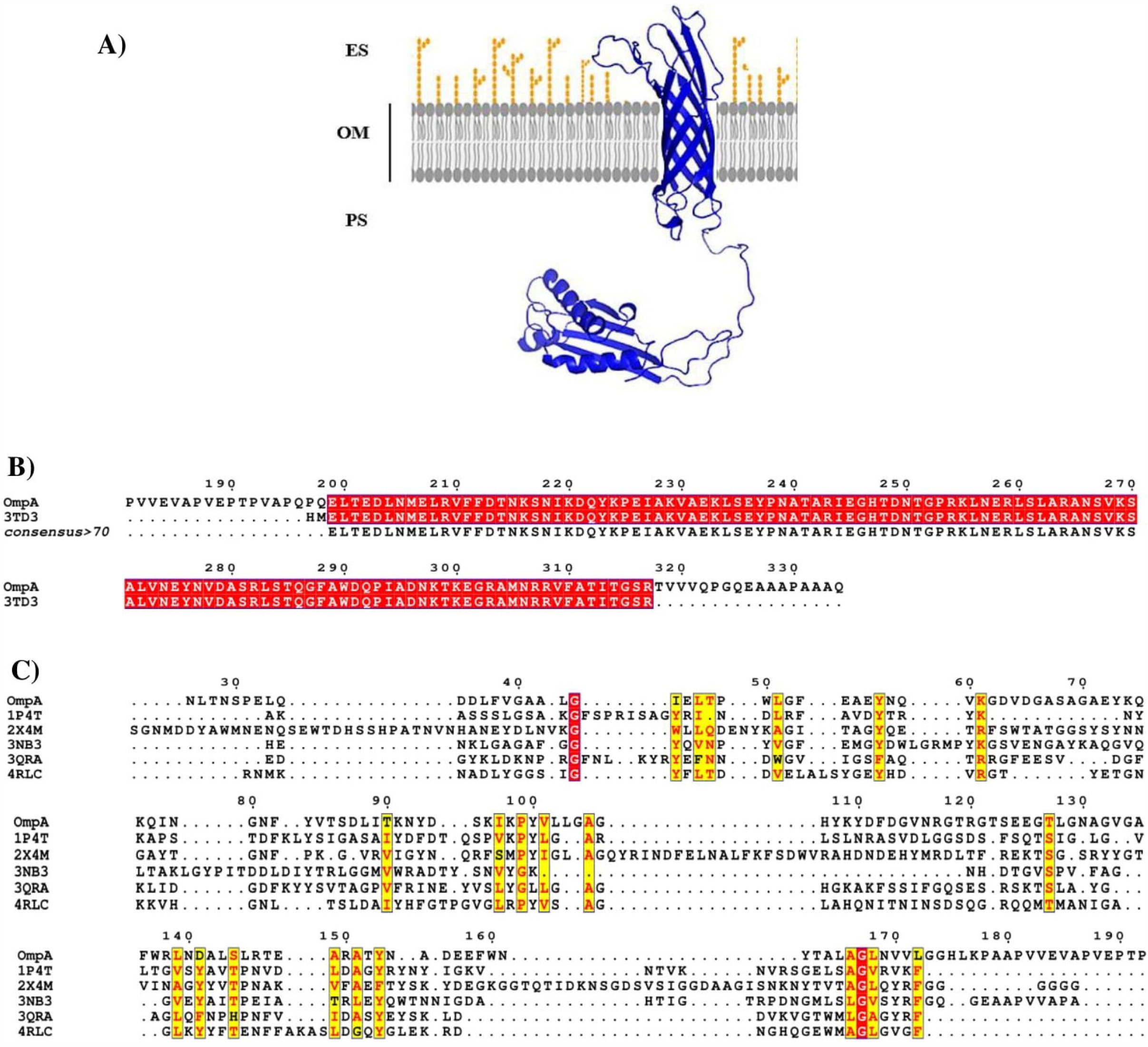
Cartoon representation of the best structures of OmpA modeled by RaptorX and refined by GalaxyRefine (**A**). Amino acid sequence alignment of OmpA and its 3 dimentional (3D) structure template for the periplasmic domain (**B**). Multiple sequence alignment of OmpA and its 3D structure templates for the β-barrel structure (**C**).

**Figure 2.**
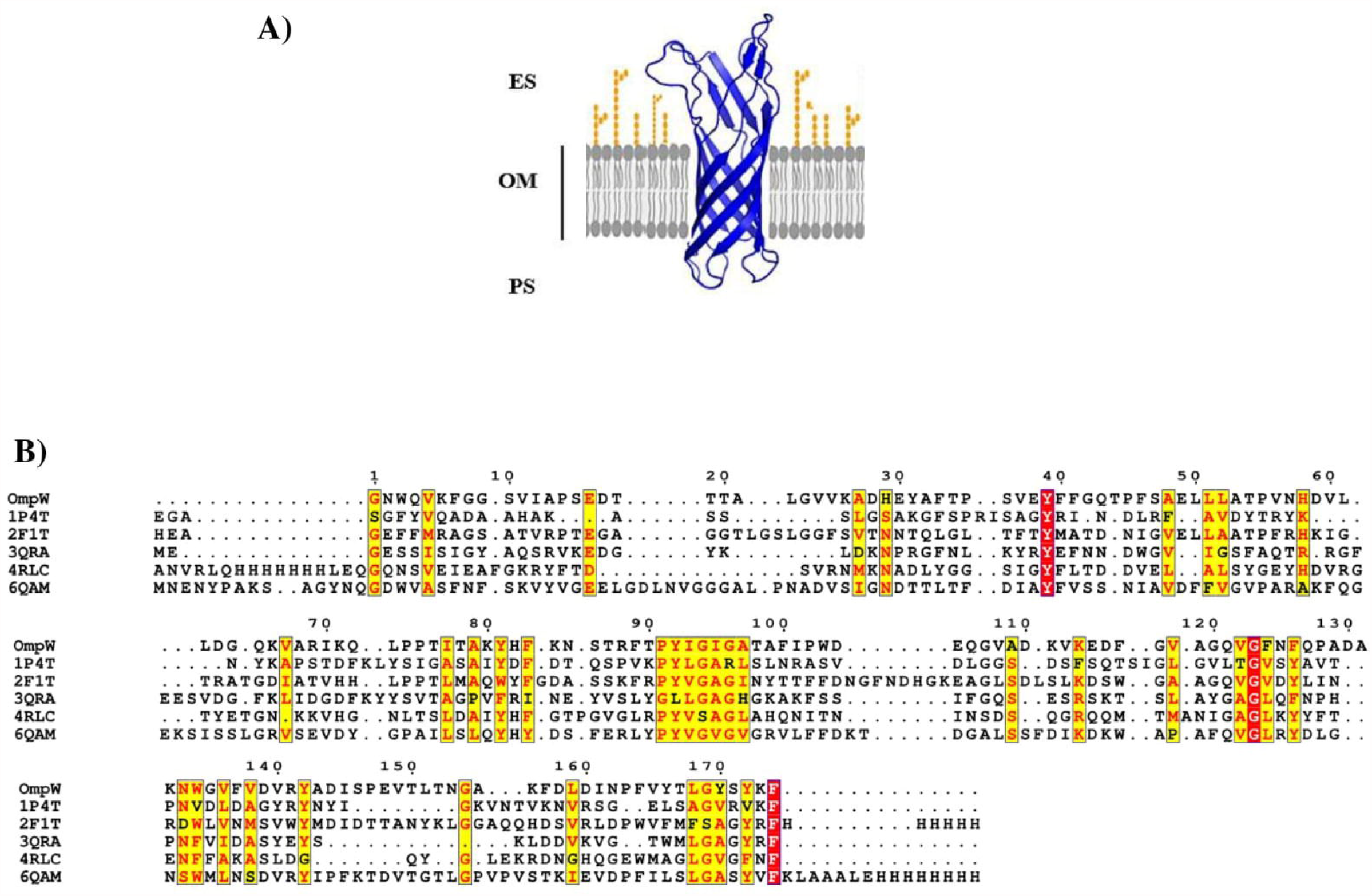
Cartoon representation of the best structures of OmpW modeled by RaptorX and refined by GalaxyRefine (**A**). Multiple sequence alignment of OmpA and its 3D structure templates (**B**).

### 3.2. Tertiary structure validation

The validity of the refined 3D modeled structures of OmpA and OmpW was checked by using RAMPAGE, ProSA-web, and ERRAT servers (Figure 3). The Ramachandran plot checked the stereochemical quality of the refined proteins. Analysis of the plot for the modeled OmpA revealed that 321 residues (96.7%) of the protein are in favored regions, which confirms that the model is characterized by stereochemical parameters of a stable structure. This is consistent with the 97.0% score predicted by the GalaxyRefine analysis. Additionally, five residues (1.5%) were predicted to be in allowed regions and six residues (1.8%) in disallowed region. The same analysis for OmpW verified that 167 residues (97.1%) are in the favored region, four residues (2.3%) are in allowed and only one residue (0.6%) is in the outlier area. The obtained result for residues of the favored region is the same as the score obtained by the GalaxyRefine analysis. These results indicate the correct geometry and three-dimensional arrangement of the models. The server ProSA-web further evaluated the overall quality of the models by providing a Z-score, as an indicative of overall model quality [43]. The reference ranges for this score change depending on the size of the protein, where the more negative Z*-*score, the better is the protein model. The values for OmpA and OmpW were -5.03 and -4.21, respectively. The Z-score of both input structures was within the range of the scores typically found for native proteins of similar sizes, which shows the modeled proteins fall within the range of X-ray solved protein structures. Further verification with PROCHECK showed a good resolution (normality) of 1.5 Å for the structures of both proteins, since most high-resolution X-ray structures have a resolution within 1.5 and 2.0 Å. The ERRAT server was used to check the statistics of non-bonded interactions between different atom types, the reliability of model proteins, and the overall quality factor; where higher scores indicate higher quality of protein structure (acceptable range= score >50). The overall quality factors were 75.44% for OmpA and 80.62% OmpW.

**Figure 3.**
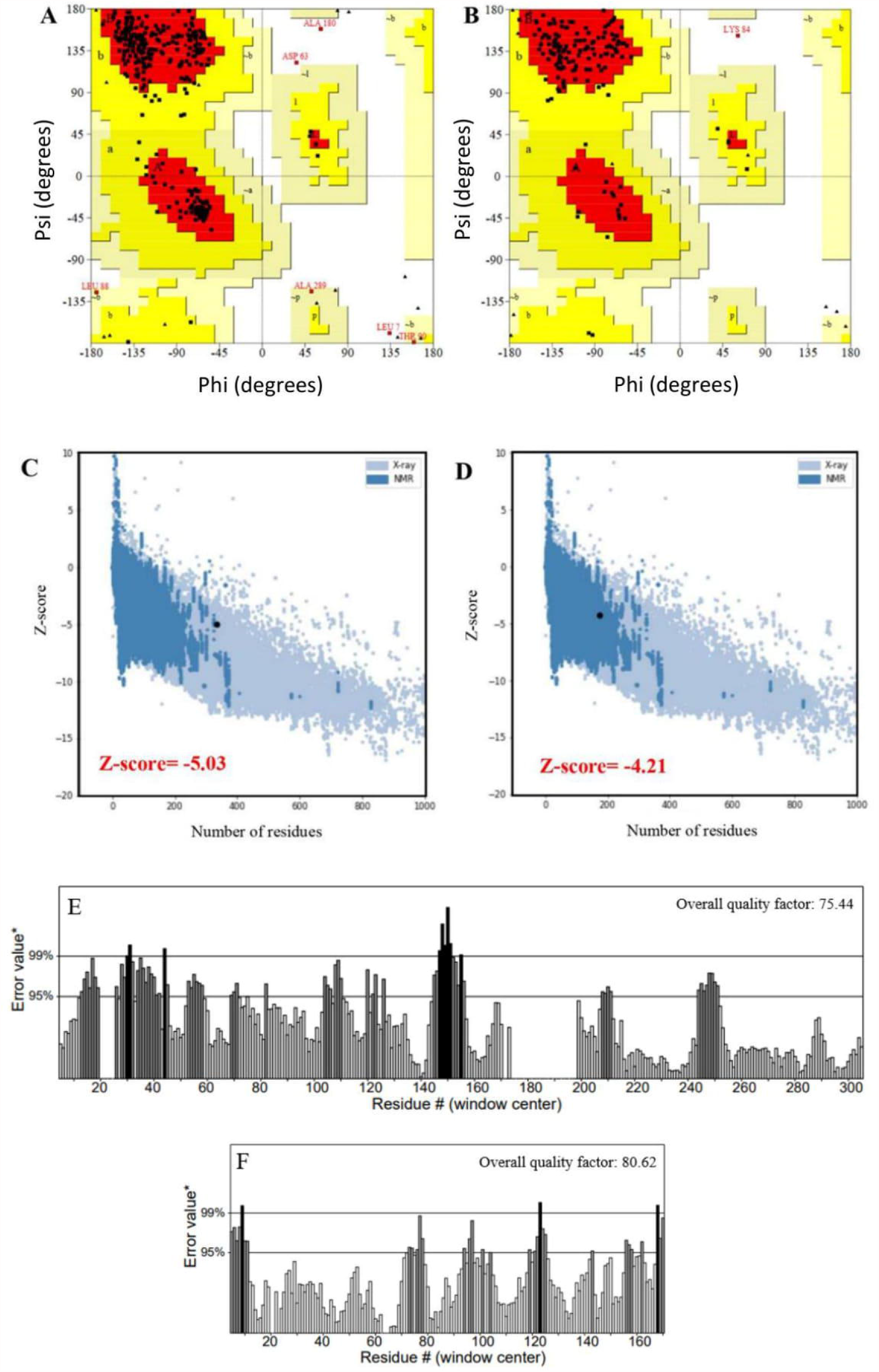
Ramachandran plot, the PROSA-web, and the ERRAT server validation of the modeled 3D structures of OmpA and OmpW. Distribution of each amino acid in the favored, allowed, and disallowed regions are shown in corresponding Ramachandran plots of OmpA (**A**), and OmpW (**B**). Ramachandran plot summary shows 96.7% residues of OmpA are in most favorable regions, 1.5% in allowed regions and 1.8% in disallowed regions. For OmpW, 97.1% residues are in most favorable regions, 2.3% in allowed regions and 0.6% in disallowed regions. Validation of OmpA (**C**) and OmpW (**D**) modeled structures using PROSA-web shows Z-score values as -5.03 and -4.21 for modeled proteins. The Overall quality factors obtained by ERRAT for OmpA (**E**) and OmpW (**F**) were 75.44% and 80.62%, respectively.

### 3.3. Pre-filtration and pharmacokinetic analysis of phytochemicals

Due to the large attrition rate of the molecules under clinical trials, and to ensure that selected compounds can be used as a drug, the putative antimicrobial phytocompounds were screened using PreADMET, SwissADME and pkCSM servers. Different filters, including Lipinski’s rule (Molecular weight ≤500 Da, logP (lipophilicity log) ≤5, number of Hydrogen Bond Donor (HBD) ≤5 and Hydrogen Bond Acceptor (HBA) ≤10, number of rotatable bonds (nRot) ≤ 10, topological polar surface area (TPSA) ≤ 150 Å2) [44], CMC like rule, MDDR like rule, lead like rule, and WDI like rule, were used to select compounds. Moreover, one important parameter for oral bioavailability is Molar Refractivity (MR) described by the Ghose filter [45]. The Ghose filter quantitatively characterizes small molecules based on computed physicochemical property profiles that include log P, molar refractivity (MR), molecular weight (MW), and the number of atoms. According to this rule, the value of MR should lie between 40 and 130 for drug-likeness.

Effective and safe drugs exhibit a good combination of pharmacodynamics (PD) and pharmacokinetics (PK) including adequate absorption, distribution, metabolism, excretion and tolerable toxicity (ADMET), which reduces the number of synthesis-evaluation cycles and more expensive late-stage failures through by excluding inappropriate drug candidates [46, 47]. Regarding this, a compound should be absorbed easily through the gastrointestinal tract so that it can be available in the systemic circulation [48]. To predict the absorption of orally administered drugs, permeability coefficient across monolayers of the human colon carcinoma cell line Caco-2 is commonly used. This permeability as log Papp (10^−6^ cm/s) rate is considered high if log Papp > 0.9 and low if < 0.9. For the achievement of an optimal clinical drug, a molecule should have high log Papp. The blood-brain barrier (BBB) and central nervous system (CNS) permeability values were also predicted for the candidates. It has been reported that those with log BB value greater than 0.3 have the potential to pass the BBB, while those with values less than -1 are poorly distributed to the brain. In addition, candidates that have log PS value > −2 are considered to penetrate the CNS, while those with values < −3 have difficulty in penetrating the CNS.

Considering the binding status of a drug with the blood proteins, the efficiency of a drug is affected by the binding efficacy of the molecule with whole blood proteins [48]. In this regard, we evaluated the fraction unbound (Fu) and steady-state volume of distribution (VDss) for the candidates. In distribution properties, VDss represents the total volume required by the drug concentration to be uniformly distributed, to give the same concentration at the target site as in the plasma. Higher values of VDss indicate that most of the drug is contained in tissues rather than in plasma (VDss (human) (log _L/kg_); low if log _L/kg_ < -0.15 and high if log _L/kg_ > 0.45).

Early *in silico* prediction of toxicity endpoints of the candidates was performed. These end points consist of the inhibition of cytochrome P450 (CYPs) monooxygenase enzymes [49]. Metabolism enzymes should metabolize a drug candidate to prevent undesirable adverse effects. In this regard, inhibition of the major human cytochrome P450 isoform of CYP2D6, which is involved in drug metabolism, as well as the renal OCT2 substrate was assessed categorically.

The factors of AMES toxicity and LD_50_ (lethal rat acute toxicity) value of the phytocompounds were investigated using the pkCSM *in silico* screening approach. Among major toxicity end-points, LD_50_ is an index determination of medicine and poison’s virulence Hepatotoxicity, skin sensitization, and inhibition of hERG potassium ion channel effects were also determined for the evaluation of drug-drug interactions (DDIs). Moreover, the solubility of the compounds was evaluated by topological methods of ESOL, Ali, and fragmental method of SILICOST-IT [48, 50].

Results showed that out of 384 investigated compounds, only 27 compounds satisfy the key parameters of these physicochemical properties (Tables S4 and S5). They fulfilled Lipinski’s rule of five, CMC like rule, lead like rule, and WDI like rule, with no violation, which are beneficial to assess *in vivo* abilities of any compounds. The ADMET profile of these candidates showed that all of the screened compounds possess high intestinal absorption. Near 63% of them showed high log Papp values. Moreover, no hepatotoxicity characterized by disrupted normal liver functions, skin sensitization, hERGI inhibition was observed for them. Our results demonstrated an acceptable prediction of physicochemical properties in comparison to Lipinski’s RO5. It was observed that selected molecules fall within most of the accepted range of the parameters of ADMET. Therefore, they were further screened against target proteins (OmpA and OmpW).

### 3.4. Protein-Ligand Interactions and Mechanisms of Binding

Previous studies have suggested that a cyclic hexapeptide (AOA-2) and dichlorophen (D6) can fight against *A. baumannii* by affecting the activity of OmpA and OmpW, respectively [24, 32, 51]. Therefore, we used these compounds as controls to evaluate the results of molecular docking. AOA-2 prevents invasion of host cells by potential binding to external loops of OmpA β-barrel structure. D6 can locate in the inner side of the OmpW β-barrel structure and interact with its residues. This can block the channel provided by this porin which consequently prevents nutrients from entering the cell.

Top-three binding cavities in the energy minimized homology models of OmpA and OmpW were predicted by the METAPOCKET target prediction tool (Figure 4). Results showed that from eight methods used by the server, only Q-SITEFINDER and POCASA algorithms failed to identify binding sites within the top predictions for both proteins. The cavity ID 03, predicted as one of OmpA binding pockets, was selected as a potential target, based on AOA-2 interaction with external loops of β-barrel structure [24]. In the case of OmpW, the cavity ID 02, containing the residues of the inner side of the β-barrel structure, was selected based on its potential interaction with D6 [32].

**Figure 4.**
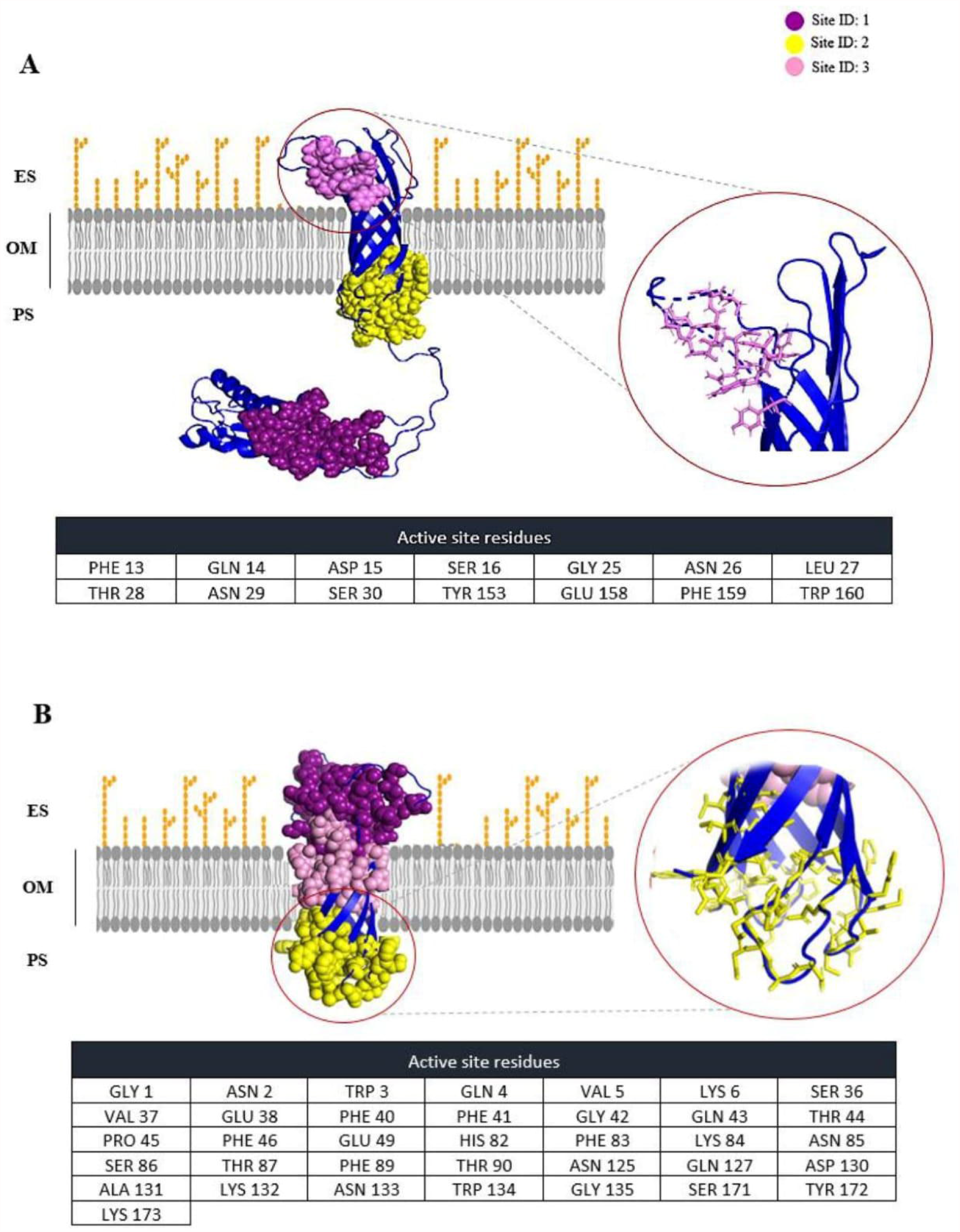
Predicted top-three potential binding pockets with corresponding active site residues for OmpA (**A**) and OmpW (**B**) by METAPOCKET 2.0. The selected potential binding pockets are emphasized by circle and their residues are shown in table.

The molecular docking results showed that isosakuranetin, aloe-emodin, and pinocembrin, with the lowest biding energy (−7.8, -7.7 and 7.5, respectively), are top three ligands to interact with the selected binding pocket of OmpA (Table 1). Although the affinity of these compounds for OmpA is lower than AOA-2, but in this regard, they can be served as stronger interacting ligands than episterol (an *A. baumannii* growth inhibitor with the ability of binding to OmpA [9]). Both isosakuranetin and pinocembrin interact with residues Gln14, Asp15, Leu27, Thr28, Ser30, Glu158, and Trp160 (Figure 5). Both compounds can interact with Asp15 by forming hydrogen bonds. In this regard, aloe-emodin is slightly different from isosakuranetin and pinocembrin. Aloe-emodin has no interaction with Ser30 and forms hydrogen bonds with Asp15 and Gln14.

**Table 1.** Molecular docking analysis of the top three compounds which have highest affinity for OmpA. The asterisks represent the compounds with the potential to inhibit OmpA according to previous studies.

| Compounds | Binding energy (kcal/mol) | RMSD | Number of H-bonds | Number of interacting residues | Interacting residues |
| --- | --- | --- | --- | --- | --- |
| **\*AOA-2** | -8.5 | 0 | 4 | 12 | Asp15, Lys23, Gly25, Asn26, Leu27, Thr28, Asn29, Ser30, Pro31, Glu157, Glu158, Phe159 |
| **\*Epiestriol** | -6.6 | 0 | 1 | 6 | Gly40, Tyr57, Asn58, Gln59, Gln75, Lys76 |
| **Isosakuranetin** | -7.8 | 0 | 1 | 7 | Gln14, Asp15, Leu27, Thr28, Ser30, Glu158, Trp160 |
| **Aloe-emodin** | -7.7 | 0 | 2 | 6 | Gln14, Asp15, Leu27, Thr28, Glu158, Trp160 |
| **Pinocembrin** | -7.5 | 0 | 1 | 7 | Gln14, Asp15, Leu27, Thr28, Ser30, Glu158, Trp160 |

**Figure 5.**
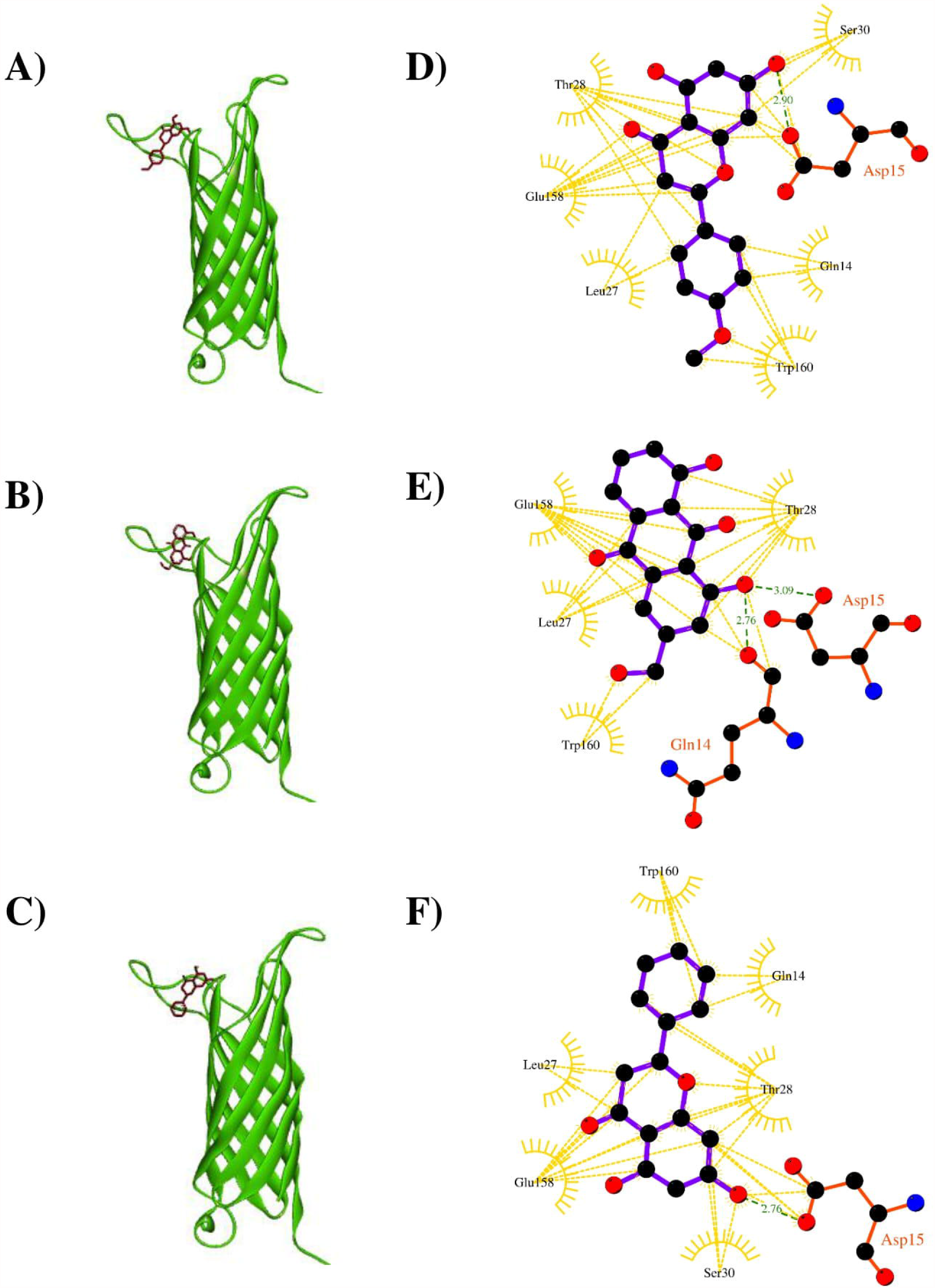
3D representation of the OmpA-ligand complexes. (**A**) represents isosakuranetin and OmpA complex, (**B**) aloe-emodin and OmpA complex, and (**C**) pinocembrin and OmpA complex. (**D**), (**E**) and (**F**) represent 2D interaction diagram of isosakuranetin, aloe-emodin and pinocembrin in the binding pockets of the target, respectively. Ligands, H-bond interacting residues and hydrophobic interacting residues are shown in purple, red, and golden, respectively. Green bonds represent hydrogen bonds and golden bonds represent hydrophobic interactions.

Compared to D6, all the above mentioned compounds possess a higher affinity for OmpW (Table 2). Isosakuranetin interacts with the highest number of residues in the selected binding pocket of OmpW. Four out of 14 residues in OmpW (Asn2, Gln42, Thr44, and Lys 84) form hydrogen bonds with isosakuranetin. Aloe-emodin and pinocembrin interact with 10 and 11 residues of OmpW, respectively. Aloe-emodin forms hydrogen bonds with 3 residues (Asn2, Gln4, and Trp134), while pinocembrin only possesses one H-bond connected to Gln4 (Figure 6).

**Table 2.** The ligand-OmpW molecular docking analysis of selected compounds. . The asterisks represent the compounds with the potential to inhibit OmpW according to previous studies.

| Compounds | Binding energy (kcal/mol) | RMSD | Number of H-bonds | Number of interacting residues | Interacting residues |
| --- | --- | --- | --- | --- | --- |
| **\*D6** | -8.4 | 0 | 1 | 12 | Asn2, Gln4, Phe40, Gly42, Gln43, Thr44, His82, Lys84, Asn85, Thr90, Asn125, Gln127 |
| **\*Epiestriol** | -6.7 | 0 | 0 | 6 | Gly1, Trp3, Tyr39, Phe40, Phe41, Ala48 |
| **Isosakuranetin** | -9.8 | 0 | 4 | 14 | Asn2, Gln4, Phe40, Gly42, Gln43, Thr44, His82, Lys84, Asn85, Thr90, Gln127, Asn133, Trp134, Ser171 |
| **Aloe-emodin** | -9.3 | 0 | 3 | 10 | Asn2, Gln4, Phe40, Gln43, Thr44, Lys84, Asn85, Thr90, Gln127, Trp134 |
| **Pinocembrin** | -10.2 | 0 | 1 | 11 | Asn2, Gln4, Phe40, Gly42, Gln43, Asn85, Asn125, Gln127, Asn133, Trp134, Ser171 |

**Figure 6.**
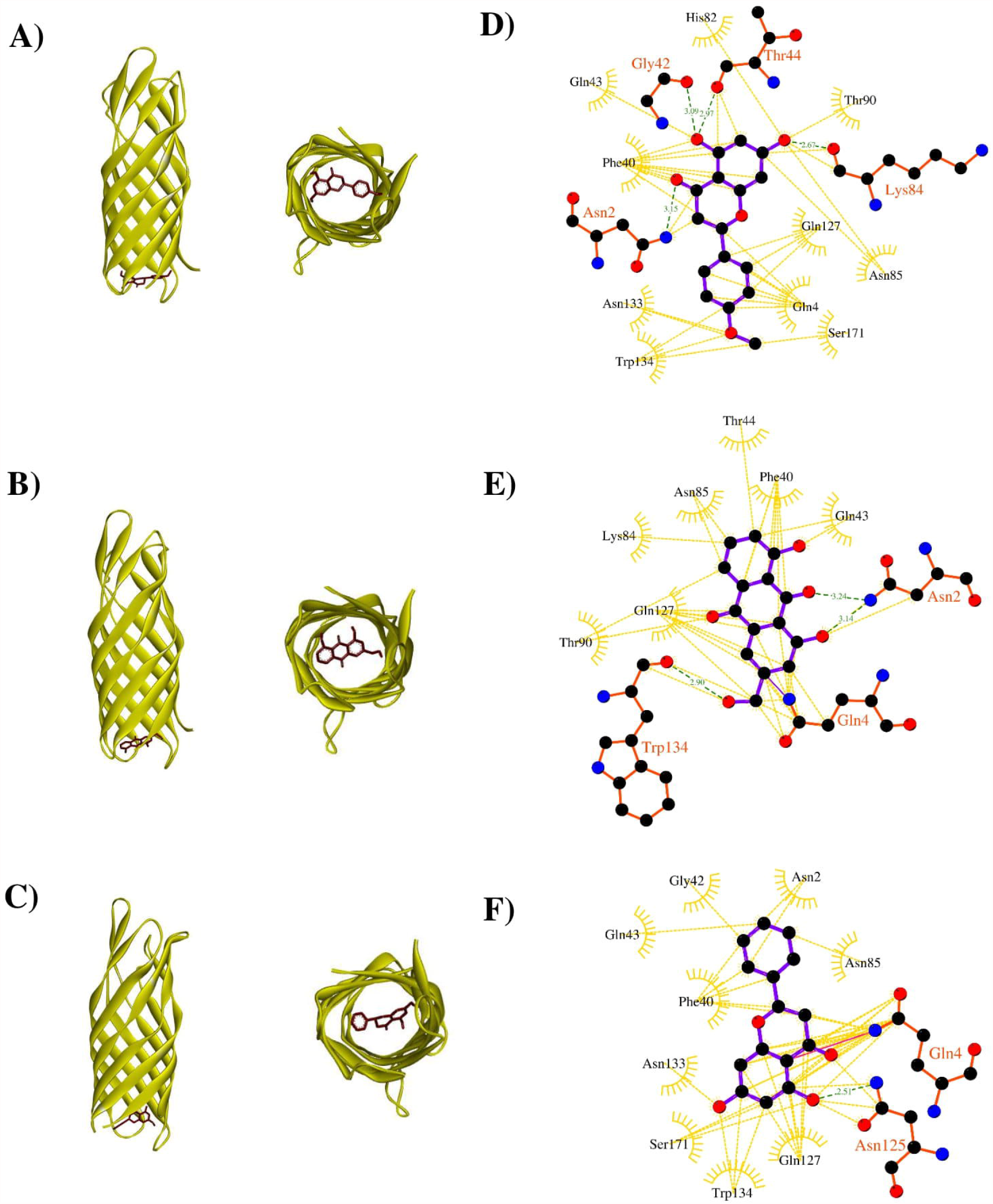
3D representation of the compound-OmpW complexes. (**A**) represents isosakuranetin and OmpW complex, (**B**) aloe-emodin and OmpW complex, and (**C**) pinocembrin and OmpW complex. (**D**), (**E**) and (**F**) represent 2D interaction diagram of isosakuranetin, aloe-emodin and pinocembrin in the binding pockets of the target, respectively. Ligands, H-bond interacting residues and hydrophobic interacting residues are shown in purple, red, and golden, respectively. Green bonds represent hydrogen bonds and golden bonds represent hydrophobic interactions.

### 3.5. Pharmacokinetic properties of selected hit phytochemicals

Physicochemical, ADMET, and safety endpoints of the selected hit phytochemicals were predicted (Table 2-4). Results showed that all of these compounds satisfy the key parameters of a drug molecule via passing in Lipinski’s RO5 (Table 3). Analyzing their ADMET profile, we observed that all of the hit inhibitors possess good gastrointestinal absorption (74.179-92.417%). Among them, the compounds isosakuranetin and pinocembrin with log Papp values of 1.1 and 1.52, respectively, showed high Caco-2 permeability. LD_50_ values of the compounds were in the range of 1.586-2.329 mol kg^-1^. None of the compounds was observed as OCT2 substrate. None of them acts as CYP2D6 substrates/inhibitors (Table 4). Additionally, the selected hits were predicted as compounds with no inhibitory effect on hERGI. They also showed no hepatotoxicity and skin sensitization activity. The high value of VDss predicted for aloe-emodin (log L/kg = 0.671 > 0.45), indicates that most amount of the drug can be contained in tissues (rather than in plasma). Since only the unbound (free) drug can diffuse between plasma and tissues, the ability of the selected compounds to interact with pharmacological target proteins such as channels was also examined. The values (Fu) were in the range of 0.022-0.226 (Table 5).

**Table 3.** The physicochemical properties of the best selected compounds with mutual inhibitory effect on both OmpA and OmpW.

| Compound name | n<sub>Rot</sub><sup>a</sup> | HBD<sup>b</sup> | HBA<sup>c</sup> | TPSA<sup>d</sup><br>(Å<sup>2</sup>) | logP<sup>e</sup> | MR<sup>f</sup> | Lipinski<br>violation | CMC-like rule | Lead-<br>like rule |
| --- | --- | --- | --- | --- | --- | --- | --- | --- | --- |
| **Isosakuranetin** | 2 | 2 | 5 | 120.92 | 2.81 | 76.04 | 0 | Qualified | Suitable |
| **Aloe-emodin** | 1 | 3 | 5 | 113.28 | 1.36 | 69.92 | 0 | Qualified | Suitable |
| **Pinocembrin** | 1 | 2 | 4 | 109.44 | 2.8 | 69.55 | 0 | Qualified | Suitable |

<sup>a</sup> Number of rotatable bonds, <sup>b</sup> Hydrogen Bond Donor, <sup>c</sup> Hydrogen Bond Acceptor, <sup>d</sup> Topological Polar Surface Area, <sup>e</sup> Lipophilicity, <sup>f</sup> Molar Refractivity.

**Table 4.** The absorption, distribution, metabolism, excretion, and toxicity (ADMET) properties of the best selected compounds with mutual inhibitory effect on both OmpA and OmpW.

| Compound name | ID | Absorption |  | Distribution |  |  | Metabolism | Excretion | Toxicity |  |
| --- | --- | --- | --- | --- | --- | --- | --- | --- | --- | --- |
|  |  | GI (%) | Water solubility (log mol/L) | Log P<sub>app</sub> | Log BB | Log PS | CYP2D6 Subs./Inh. | OCT2 Subs | AMES Toxicity | LD50 (mol/kg) |
| Isosakuranetin | 160481 | 92.37 | -3.092 | 1.1 | 0.194 | -2.17 | No/No | No | No | 1.843 |
| Aloe-emodin | 10207 | 74.179 | -3.104 | -0.233 | -0.729 | -2.466 | No/No | No | Yes | 2.329 |
| Pinocembrin | 68071 | 92.417 | -3.538 | 1.152 | 0.42 | -2.047 | No/No | No | No | 1.586 |

**Table 5.**
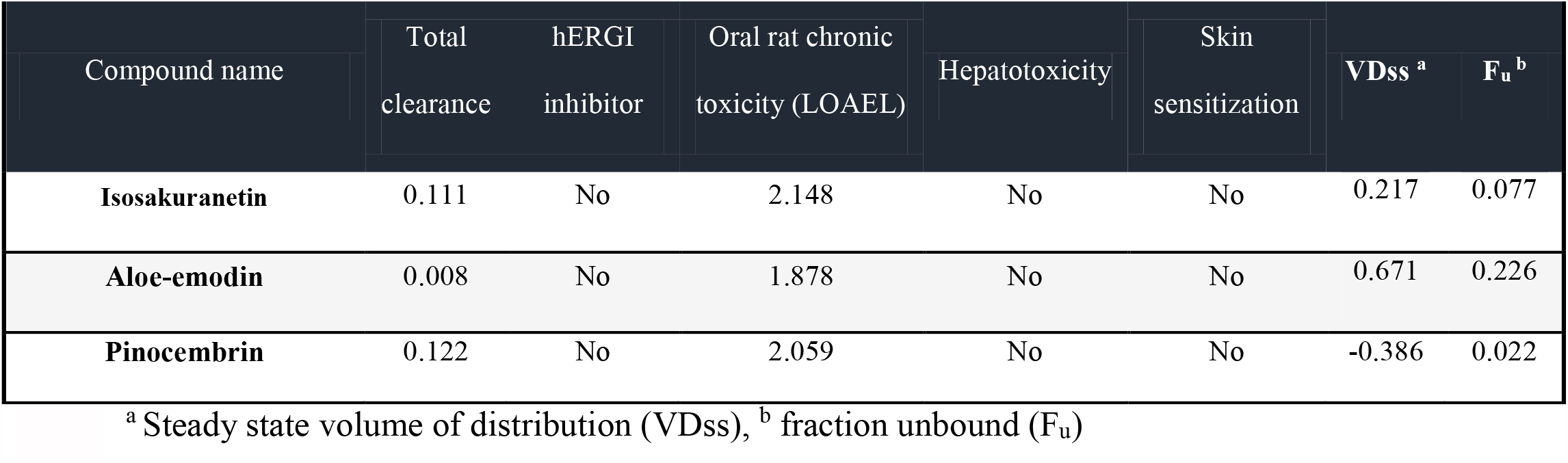
The computed safety endpoints of the best selected compounds with mutual inhibitory effect on both OmpA and OmpW.

Apart from efficacy and toxicity, poor pharmacokinetics and bioavailability are among important factors of drug development failures. Among these, gastrointestinal absorption and brain access are crucial pharmacokinetic features, which need to be estimated in drug discovery processes. Permeation from the blood–brain barrier (BBB), as a physical and biochemical shield protecting the brain, is the basic of the distribution of central-acting molecules. Moreover, regarding the preference of oral bioavailability of a drug in drug administration, Human Intestinal Absorption (HIA) is an important roadblock in drug research. The accurate predictive model of Brain Or IntestinaL EstimateD permeation method (BOILED-Egg) was used to predict these two important behaviors for selected phytocompounds (Figure 7). The model calculates the lipophilicity and polarity of small molecules. It allows for the evaluation of HIA as a function of the position of the small molecules in the WLOGP-versus-TPSA referential. The white region of the “BOILED-Egg” is for a high probability of passive absorption of a compound by the gastrointestinal tract, and the yellow region (yolk) is for a high probability of brain penetration. Yolk and white areas are not mutually exclusive. In addition, the points are colored in blue if a compound predicted as actively effluxed by P-glycoprotein (PGP+) and in red if a compound predicted as non-substrate of P-gp (PGP−). Our results showed that among the selected compounds, isosakuranetin and pinocembrin were predicted to permeate through BBB by passive diffusion. According to the prediction, aloe-emodin can be passively absorbed by the gastrointestinal tract. The assessment of P-gp efflux for the selected compounds showed all of the selected compounds are not substrates of P-gp. Indeed, P-glycoprotein is a multidrug transporter, which is apically expressed in the gastrointestinal tract, liver, kidney and brain endothelium [52]. Consequently, P-glycoprotein plays an important role in the oral bioavailability, CNS distribution biliary and renal elimination of drugs which are substrates of this transporter.

**Figure 7.**
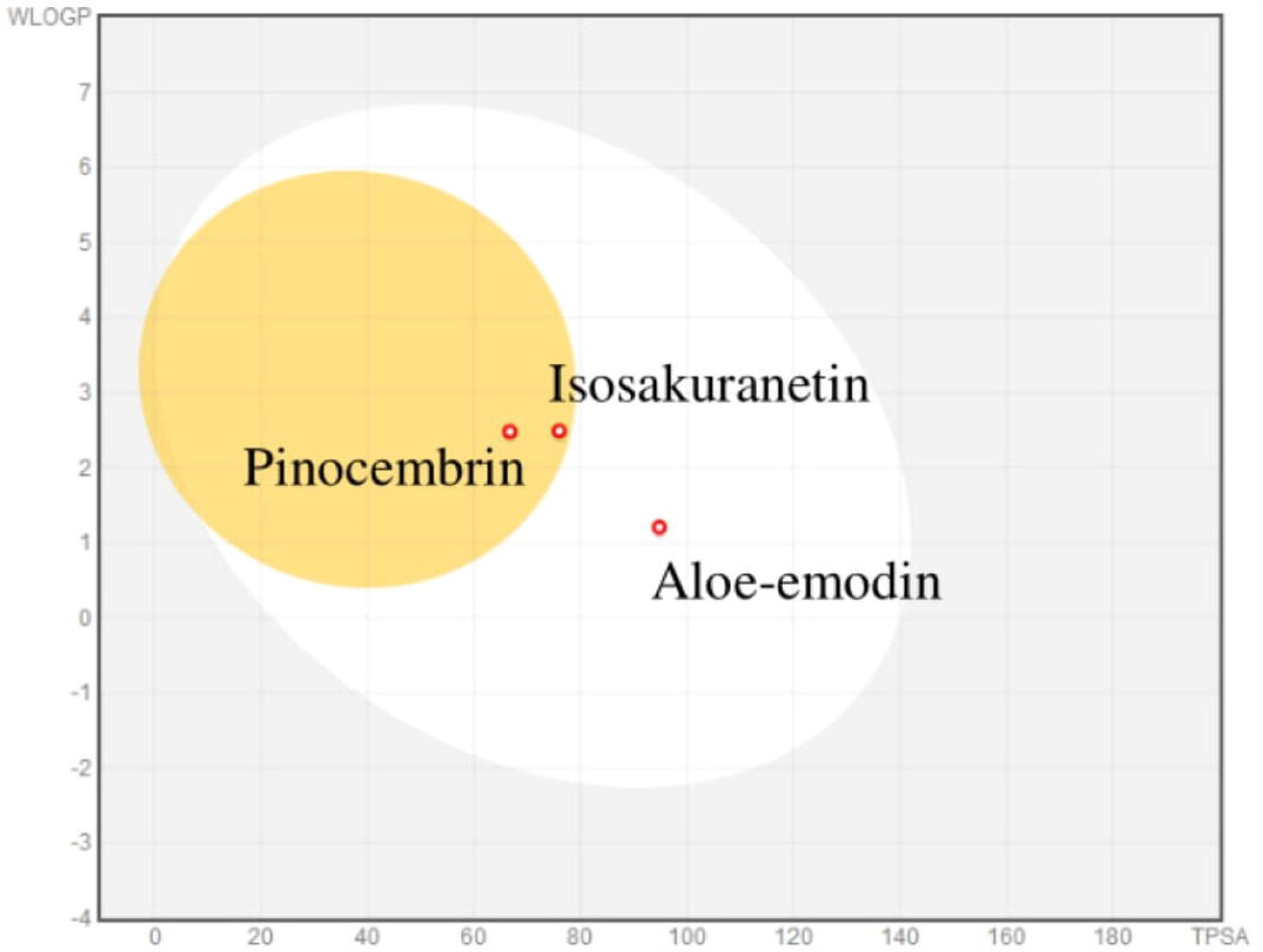
The BOILED-Egg model of the selected best potential inhibitors of OmpA and OmpW. The compounds located in the yellow region were predicted to be passively penetrate from BBB. The compound in the white region is predicted to be passively absorbed by gastrointestinal tract. The molecules predicted to be effluated from the central nervous system by p-glycoproteins are shown by blue dots. Those which do not effluated from the central nervous system by P-glycoproteins are shown by red dots.

HIA and BBB are dependent on the water solubility and lipophilicity of the drug. Two topological methods of the ESOL [53] and Ali [54] are included in SwissADME to predict water solubility. SwissADME third predictor for solubility was developed by SILICOS-IT. All predicted values are the decimal logarithm of the molar solubility in water (log S) (Table 5). Three predictors classified aloe-emodin and pinocembrin as soluble molecules. Isosakuranetin is predicted to be soluble adopted by Esol predictor, but moderately soluble by Ali and SILICOS-IT predictors. According to Log P, isosakuranetin (Log p= 2.81) and pinocembrin (Log p= 2.8) are more lipophilic than aloe-emodin (Log p= 1.36) (Table 6).

**Table 6.** The water solubility evaluation of the best selected compounds with mutual inhibitory effect on both OmpA and OmpW.

| Molecule | ESOL LogS | Ali LogS | SILICOS-IT LogS | ESOL Class | Ali Class | SILICOS-IT Class |
| --- | --- | --- | --- | --- | --- | --- |
| **Isosakuranethin** | -3.7 | -4.1 | -4.12 | Soluble | Moderately soluble | Moderately soluble |
| **Aloe-emodin** | -3.04 | -3.43 | -3.92 | Soluble | Soluble | Soluble |
| **Pinocembrin** | -3.64 | -3.94 | -4.00 | Soluble | Soluble | Soluble |

### 3.6 Molecular dynamics simulation analysis

Given the results, it seems isosakuranetin and aloe-emodin can be served as suitable ligands for both OmpA and OmpW to inhibit their targeted functions. Therefore, both of the compounds were selected to evaluate their molecular docking results using MD simulation (Figure 8). Backbone deviation of complexes were plotted as a function of time. The results showed stability of complexes. The aloe emodin-OmpA complex showed a RSMD value ranging from 0.1 to 0.65 nm during simulation. RMSD value for the isosakuranetin-OmpA complex were in the range of 0.1 to 0.65 nm. Both the isosakuranetin- and aloe emodin-OmpW complex demonstrated a similar value, ranging from 0.1 to 0.4 nm. The flexibility of backbone was evaluated using RMSF plot. The higher RMSF value means the higher flexibility, whereas the lower value represents the lower mobility during simulation. In general, β-strand residues showed less fluctuations than loops. Compared to the aloe emodin-OmpA complex, the isosakuranetin-OmpA complex demonstrated lower fluctuations. In regard to the OmpW-ligand complexes, the isosakuranetin-OmpW complex showed slightly higher fluctuations than the aloe emodin-OmpW complex.

**Figure 8.**
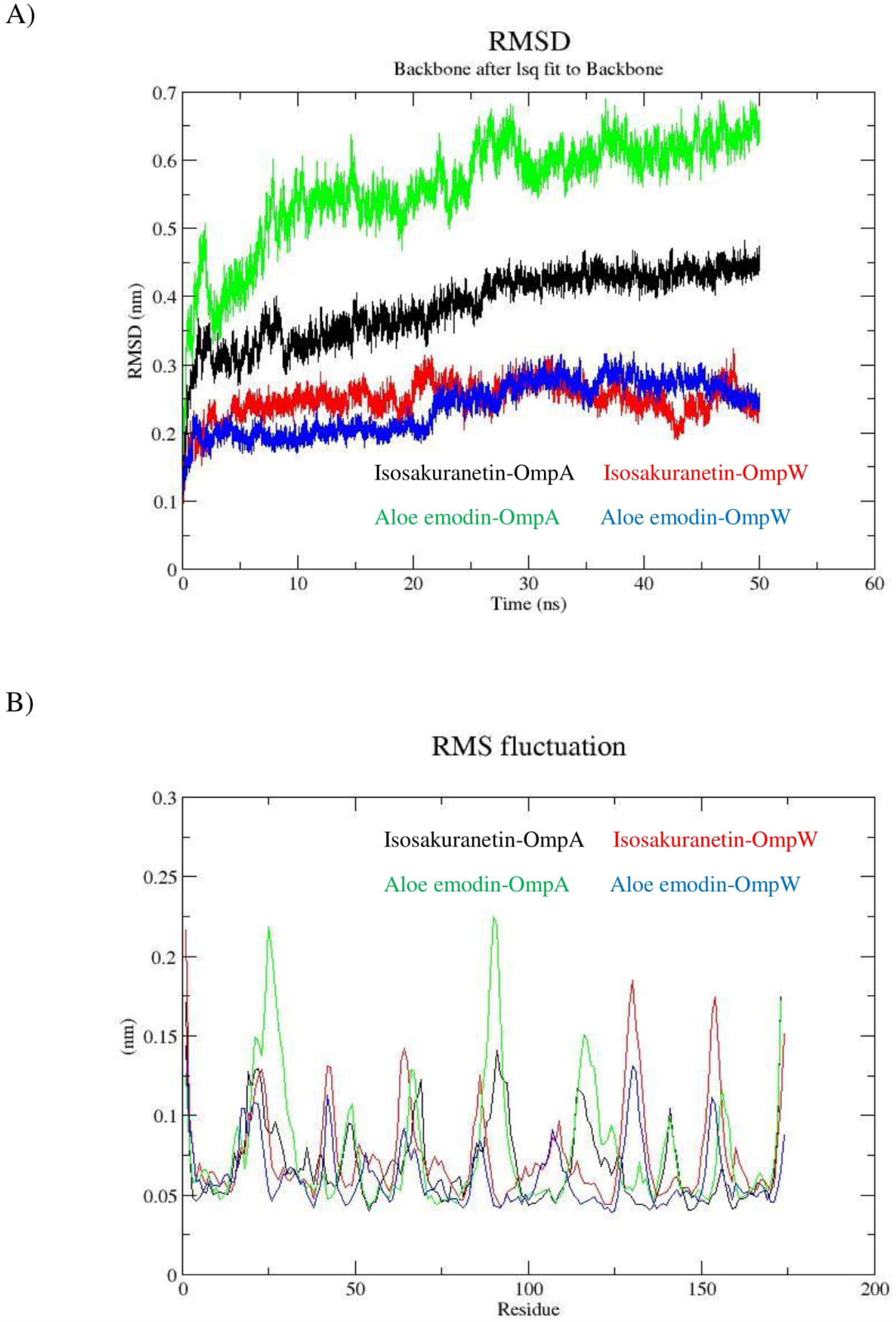
Detection of the stability of the protein-ligand complexes. (**A**) RMSD of the protein-ligand complexes. (**B**) Root mean square fluctuation (RMSF) plots of the protein-ligand complexes. Isosakuranetin bounded to OmpA (black) and bounded to OmpW (red). Aloe-emodin bounded to OmpA (green) and bounded to OmpW (blue).

**Figure 9.**
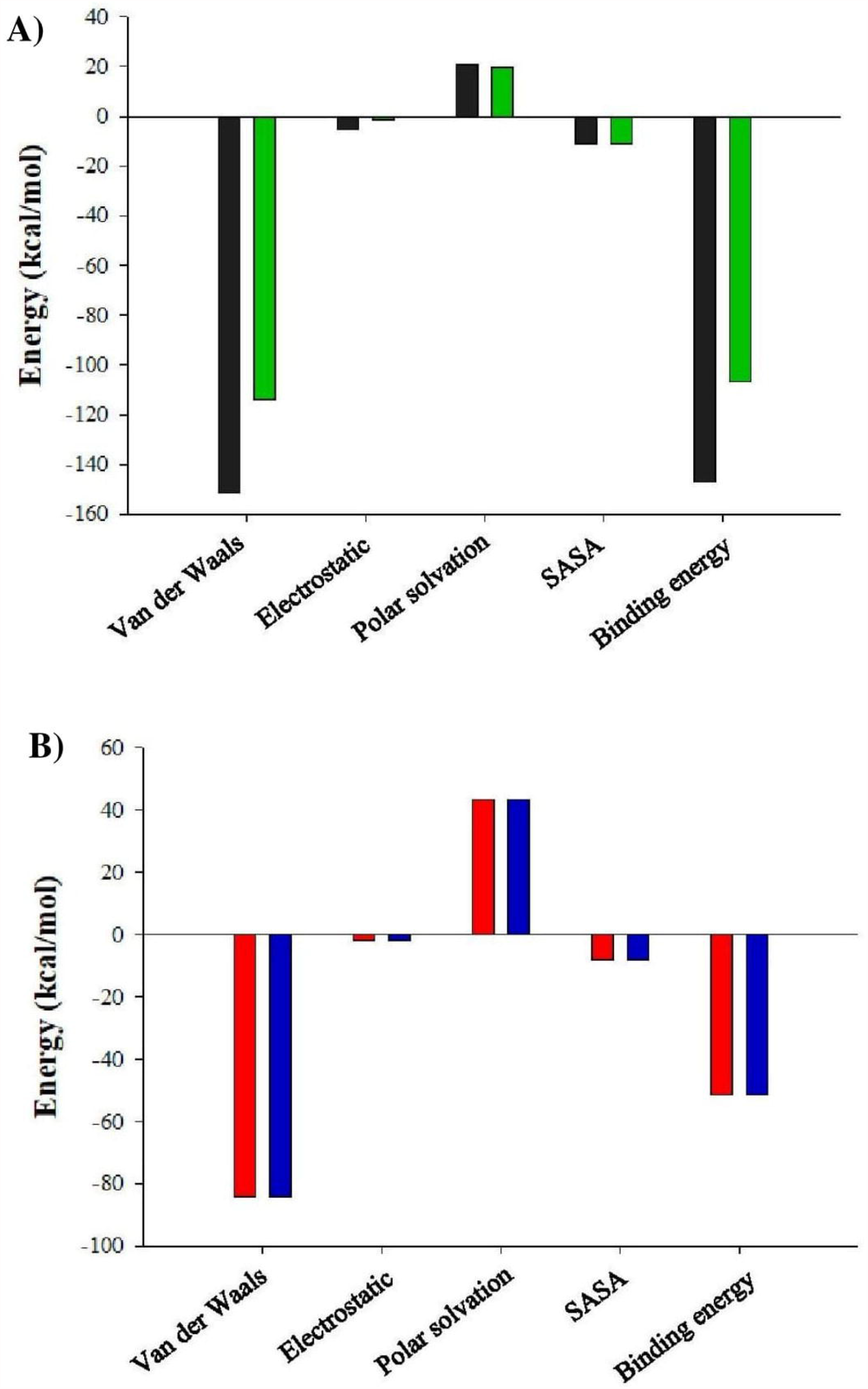
(**A**) Binding energies (including its components) of isosakuranetin-OmpA and aloe emodin-OmpA (black and green bars, respectively). (**B**) Binding energies (including its components) of isosakuranetin-OmpW and aloe emodin-OmpW (red and blue bars, respectively).

We performed MM/PBSA analysis to evaluate free-binding energy of the ligand-protein complexes. Compared to the aloe-emodin, isosakuranetin interacts with OmpA with larger Van der Waals and electrostatic energy value. The binding energies for isosakuranetin and aloe-emodin, in the form of OmpA-ligand complex, were -147.097 and -106.871 (kcal/mol), respectively. According to OmpW-ligand complex, both isosakuranetin and aloe-emodin form complexes with the same binding energy (−51.49 kcal/mol). Overall, although the results show that both compounds are suitable for the purpose of our study, isosakuranetin appears to establish more stable interactions with both OmpA and OmpW.

## 4. Conclusion

The global antimicrobial resistance problem raises the necessity of novel approaches for the development of new drugs against emerging superbugs. A major contributor to this resistance is the outer membrane of Gram-negative bacteria with outer membrane proteins (OMPs) as promising drug targets due to their extensive functions. Therefore, any molecule which has the potential to disrupt the normal activity of these porins can help the host defense system overcoming the pathogens. Our results suggest three out of 384 phytocompounds (isosakuranetin, aloe-emodin, and pinocembrin), can control *A. baumannii* infection by targeting the function of two proteins, OmpA, based on its important roles in biofilm formation and bacterial adhesion to host cells, and OmpW, based on its important role in the transport of essential nutrients. Also, these compounds showed promising pharmacokinetic features as potential drugs with possible anti-*A*.*baumannii* activities. Amongst them, isosakuranetin was introduced as a top inhibitor for the targeted functions of OmpA and OmpW. Therefore, this biomolecule can be further characterized experimentally to corroborate its inhibitory activity. In addition, the skeletons of these active compounds could be adopted as starting potential scaffolds for the design of future anti-*A*.*baumannii* drugs.

## Data accessibility

The data supporting the results in this article can be accessed at the Dryad Digital Repository: https://doi.org/10.5061/dryad.k6djh9w6f https://datadryad.org/stash/share/joYqZpWMp7ssFCmDdVSVy3lMdnYlBQ6qyK2r_bJaubg

