## Supplementary material for "Screening of anti-*Acinetobacter baumannii* phytochemicals, based on the potential inhibitory effect on OmpA and OmpW functions": Supplementary materials.docx

**Table S1.** Function and description of the web servers and tools used in the study.

| Tool | Function | Description | URL |
| --- | --- | --- | --- |
| NCBI | Development of new information technologies to aid in the understanding of fundamental molecular and genetic processes that control health and disease. | The National Center for Biotechnology Information advances science and health by providing access to biomedical and genomic information | <https://www.ncbi.nlm.nih.gov/> |
| SignalP 4.1 | server predicts the presence and location of signal peptide cleavage sites in amino acid sequences from different organisms | The method incorporates a prediction of cleavage sites and a signal peptide/non-signal peptide prediction based on a combination of several artificial neural networks. | <http://www.cbs.dtu.dk/services/SignalP-4.1/> |
| LipoP 1.0 | Predictions of signal peptides | The server produces predictions of lipoproteins and discriminates between lipoprotein signal peptides, other signal peptides and n-terminal membrane helices in Gram-negative bacteria. | <http://www.cbs.dtu.dk/services/LipoP> |
| Raptor X | Tertiary Structure Prediction | RaptorX is developed by Xu group, excelling at secondary, tertiary and contact prediction for protein sequences without close homologs in the Protein Data Bank (PDB). | <http://raptorx.uchicago.edu> |
| Galaxy Refine Server | Refinement of Tertiary Structure | A web server to improve the structure of protein models. It is particularly effective in improving the quality of local structures as demonstrated by the CASP refinement category. | <http://galaxy.seoklab.org/> |
| RAMPAGE SERVER | Validation of Tertiary Structure | Rampage server is used for the validation of 3d structure modeled by plotting Ramachandran plot | <http://mordred.bioc.cam.ac.uk/~rapper/rampage.php> |
| ProSA-web | ProSA calculates an overall quality score for a specific input structure. | ProSA-web provides an easy-to-use interface to the program ProSA (Sippl 1993) which is frequently employed in protein structure validation | <https://prosa.services.came.sbg.ac.at/prosa.php> |
| ERRAT | Analyzes the statistics of non-bonded interactions between different atom types | The server plots the value of the error function versus position of a 9-residue sliding window, calculated by a comparison with statistics from highly refined structures. | <http://services.mbi.ucla.edu/ERRAT/> |
| METAPOCKET | a meta server to identify ligand binding sites on protein surface | It is a consensus method, in which the predicted binding sites from eight methods:  LIGSITE, PASS, Q-SiteFinder, SURNET, Fpocket, GHECOM, ConCavity and POCASA are combined together to improve the prediction success rate. | <https://projects.biotec.tu-dresden.de/metapocket/index.php> |
| SwissADME | physicochemical descriptors calculations | The server computes physicochemical descriptors as well as to predict ADME parameters, pharmacokinetic properties, druglike nature and medicinal chemistry friendliness of one or multiple small molecules to support drug discovery. | <http://www.swissadme.ch/> |
| PubChem | PubChem is a database of chemical molecules and their activities against biological assays. | The system is maintained by the National Center for Biotechnology Information, a component of the National Library of Medicine, which is part of the United States National Institutes of Health. | https://pubchem.ncbi.nlm.nih.gov/ |
| pkCSM | Prediction of small-molecule pharmacokinetic properties using graph-based signatures | A novel approach to the prediction of pharmacokinetic properties, called pkCSM, which relies on graph-based signatures | <http://biosig.unimelb.edu.au/pkcsm/> |
| PyMOL | a user-sponsored molecular visualization system | An open-source molecular visualization system, which can produce high-quality 3D images of small molecules and biological macromolecules. | <https://pymol.org> |
| AutoDock Vina | a molecular modeling simulation software. It is especially effective for protein-ligand docking. | AutoDock Vina is a successor of AutoDock, significantly improved in terms of accuracy and performance. | http://vina.scripps.edu/ |
| ChemDraw | ChemDraw is a [molecule editor](https://en.wikipedia.org/wiki/Molecule_editor) | The software possess different features. | https://perkinelmerinformatics.com/products/research/chemdraw/ |
| MEGA X | Molecular Evolutionary Genetics Analysis is computer software for conducting statistical analysis of molecular evolution and for constructing phylogenetic trees. | It includes many sophisticated methods and tools for phylogenomics and phylomedicine. | https://www.megasoftware.net/ |
| GROMACS | a molecular dynamics package mainly designed for simulations of proteins, lipids, and nucleic acids. | Written in \| [C++](https://en.wikipedia.org/wiki/C%2B%2B), [C](https://en.wikipedia.org/wiki/C_(programming_language)), [CUDA](https://en.wikipedia.org/wiki/CUDA), [OpenCL](https://en.wikipedia.org/wiki/OpenCL), SYCL | https://www.gromacs.org/ |

**Table S2.** Summary of phytochemicals used in this work

| **No.** | **Compound name** | **PubChem CID** | **Compound structure** | **Physicochemical Properties** | |
| --- | --- | --- | --- | --- | --- |
|  |  |  |  | **Formula** | **Mw**  **(g/mol)** |
| 1 | Indole | 798 | 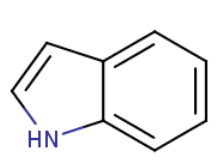 | C_8_H_7_N | 117.15 |
| 2 | 3-Indolylacetonitrile | 351795 | 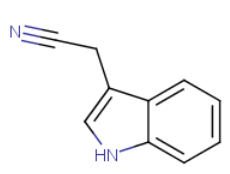 | C_10_H_8_N_2_ | 156.18 |
| 3 | Indole-3-carbinol | 3712 | 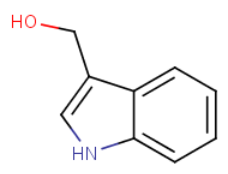 | C_9_H_9_NO | 147.17 |
| 4 | Indole-3-acetaldehyde | 800 | 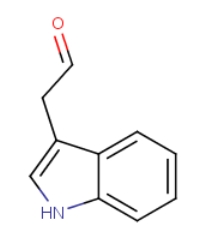 | C_10_H_9_NO | 159.18 |
| 5 | Indole-3-carboxyaldehyde | 10256 | 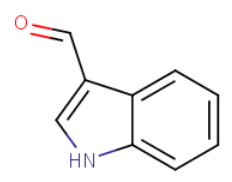 | C_9_H_7_NO | 145.16 |
| 6 | Indole-3-acetamide | 397 | 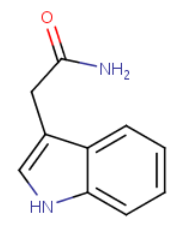 | C_10_H_10_N_2_O | 174.20 |
| 7 | Indirubin | 10177 | 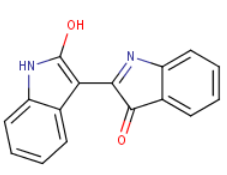 | C_6_H_10_N_2_O_2_ | 262.26 |
| 8 | Isatin | 7054 | 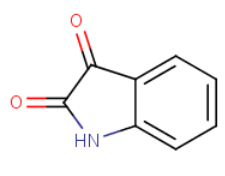 | C_8_H_5_NO_2_ | 147.13 |
| 9 | Reserpine | 5770 | 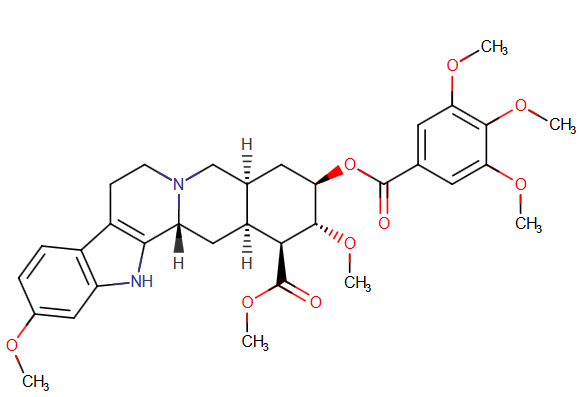 | C_33_H_40_N_2_O_9_ | 608.68 |
| 10 | Deoxynojirimycin | 29435 | 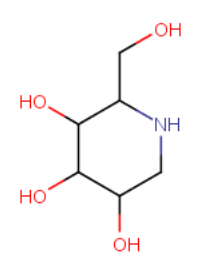 | C_6_H_13_NO_4_ | 163.17 |
| 11 | Piperine | 638024 | 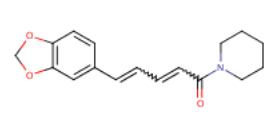 | C_17_H_19_NO_3_ | 285.34 |
| 12 | Tomatidine | 65576 | 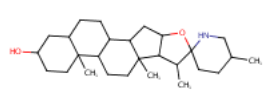 | C_27_H_45_NO_2_ | 415.65 |
| 13 | Kinurenic acid | 3845 | 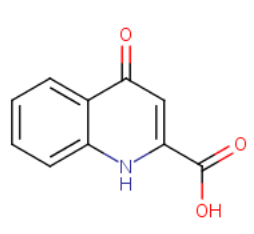 | C_10_H_7_NO_3_ | 189.17 |
| 14 | Berberine | 2353 | 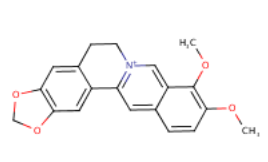 | C_20_H_18_NO_4_ | 336.36 |
| 15 | Sanguinarine | 5154 | 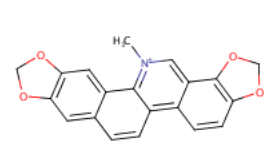 | C_20_H_14_NO_4_ | 332.33 |
| 16 | Chelerythrine | 2703 | 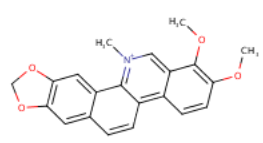 | C_21_H_18_NO_4_ | 348.37 |
| 17 | Palmatine | 19009 | 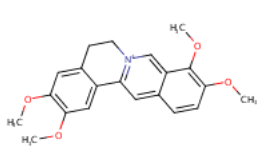 | C_21_H_22_NO_4_ | 352.40 |
| 18 | Coptisine | 72322 | 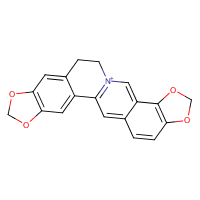 | C_19_H_14_NO_4_ | 320.32 |
| 19 | Pseudodehydrocorydaline | - | 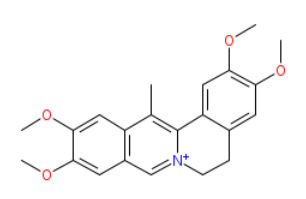 | C_22_H_24_NO_4_ | 366.430 |
| 20 | Jatrorrhizine | 72323 | 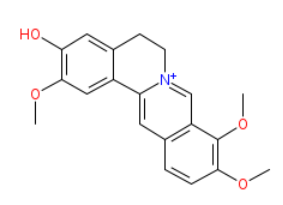 | C_20_H_20_NO_4_ | 338.377 |
| 21 | Dehydrocorybulbine | 5316439 | 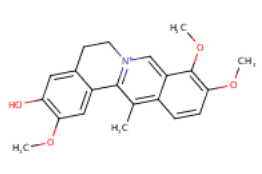 | C_21_H_22_NO_4_ | 352.4 |
| 22 | Pseudocoptisine | 15520811 | 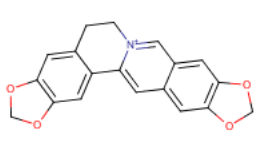 | C_19_H_14_NO_4_ | 320.32 |
| 23 | Anisodamine | 183088 | 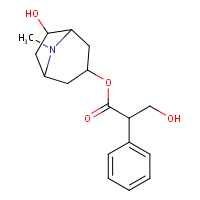 | C_17_H_23_NO_4_ | 305.4 |
| 24 | Caffeine | 2519 | 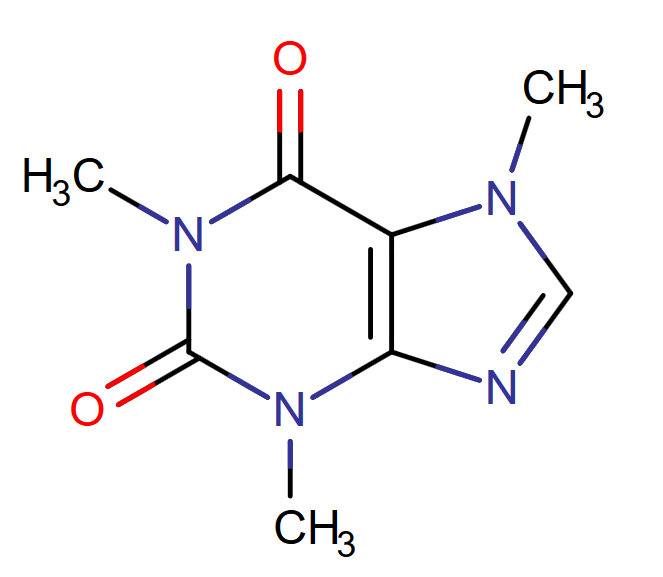 | C_8_H_10_N_4_O_2_ | 194.19 |
| 25 | Capsaicin | 1548943 | 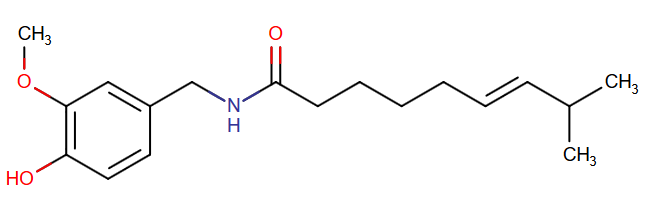 | C_18_H_27_NO_3_ | 305.41 |
| 26 | 11-Methyldodecanoic acid (iso-C 13:0) | 33002 | 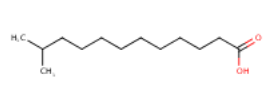 | C_13_H_26_O_2_ | 214.34 |
| 27 | Lauric acid | 3893 | 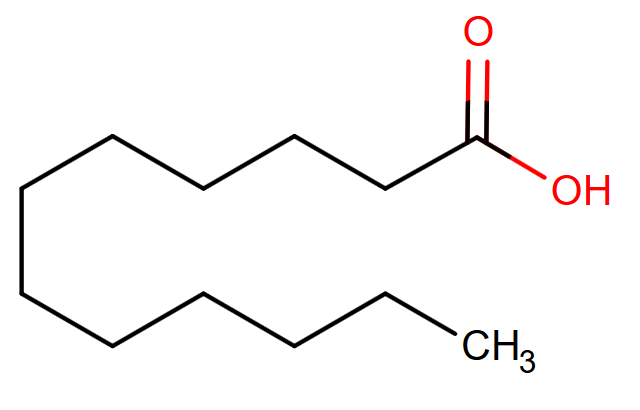 | C_12_H_24_O_2_ | 200.32 |
| 28 | Myristic acid | 11005 | 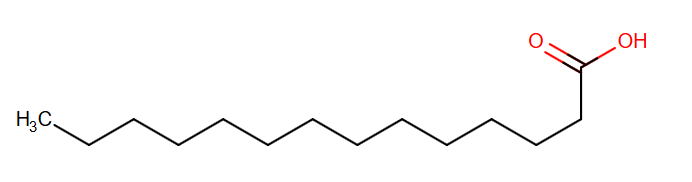 | C_14_H_28_O_2_ | 228.37 |
| 29 | Palmitic acid | 985 | 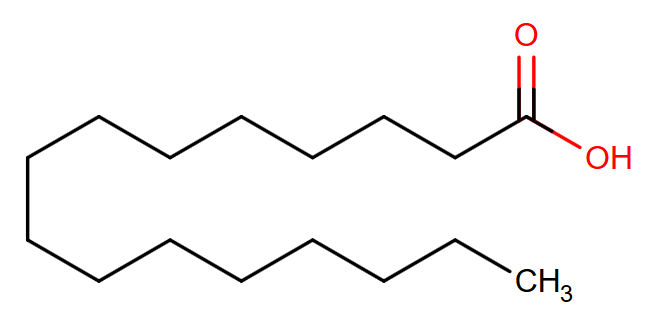 | C_16_H_32_O_2_ | 256.42 |
| 30 | Stearic acid | 5281 | 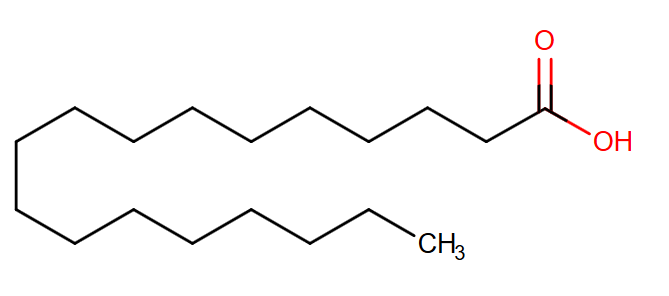 | C_18_H_36_O_2_ | 284.48 |
| 31 | Vaccenic acid (cis-11-octadecenoic acid) | 5282761 | 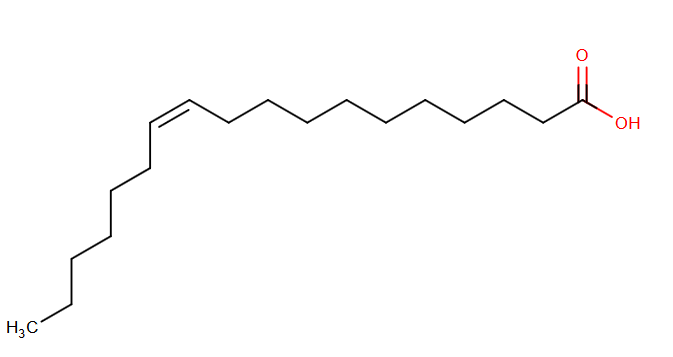 | C_18_H_34_O_2_ | 282.46 |
| 32 | Oleic acid  (cis-9-octadecenoic acid) | 445639 | 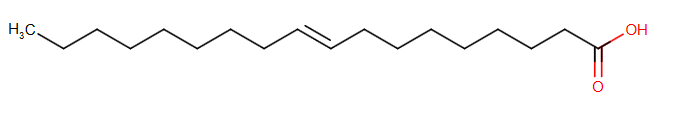 | C_18_H_34_O_2_ | 282.46 |
| 33 | Linoleic acid  (cis-9, cis-12-octadecadienoic acid) | 5280450 | 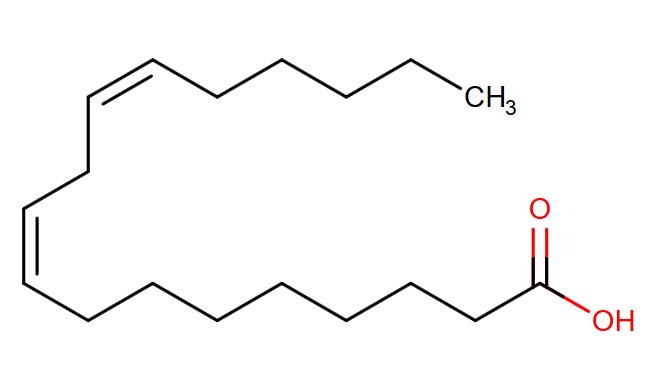 | C_18_H_32_O_2_ | 280.45 |
| 34 | Allyl isothiocyanate | 5971 | 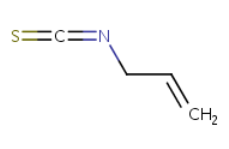 | C_4_H_5_NS | 99.15 |
| 35 | Benzyl isothiocyanate | 2346 | 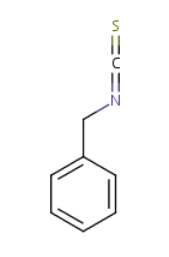 | C_8_H_7_N_S_ | 149.21 |
| 36 | 2-Phenylethyl-isothiocyanate | 16741 | 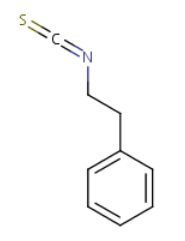 | C_9_H_9_NS | 163.24 |
| 37 | Allicin | 65036 | 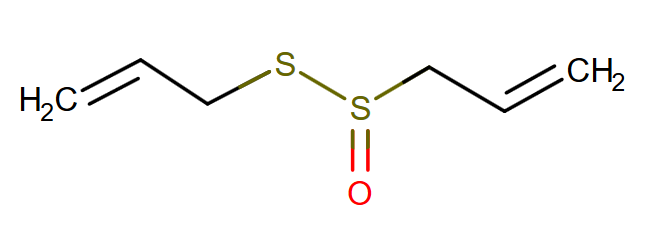 | C_6_H_10_OS_2_ | 162.27 |
| 38 | Ajoene | 5386591 | 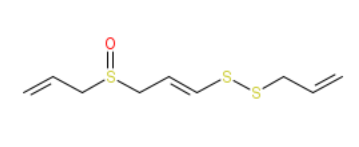 | C_9_H_14_OS_3_ | 234.40 |
| 39 | Iberin | 10455 | 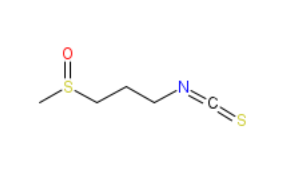 | C_5_H_9_NOS_2_ | 163.26 |
| 40 | Diallyl trisulphide | 16315 | 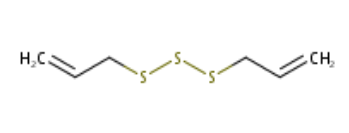 | C_6_H_10_S_3_ | 178.34 |
| 41 | Sulforaphane | 5350 | 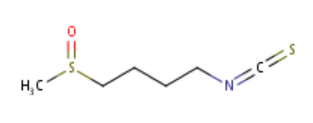 | C_6_H_11_NOS_2_ | 177.29 |
| 42 | Erucin | 78160 | 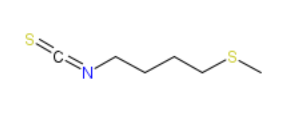 | C_6_H_11_NS_2_ | 161.29 |
| 43 | Methyl styryl sulfone | 138463 | 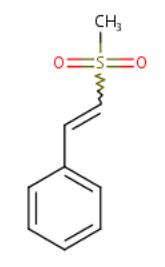 | C_9_H_10_O_2_S | 182.24 |
| 44 | Zosteric acid | 6070438 | 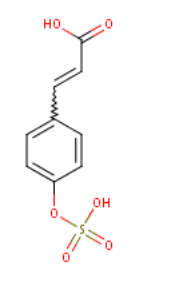 | C_9_H_8_O_6_S | 244.22 |
| 45 | Trans-2-Hexen-1-al | 10460 | 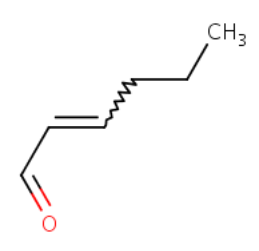 | C_6_H_10_O | 98.14 |
| 46 | Trans-2-Heptenal | 5283316 | 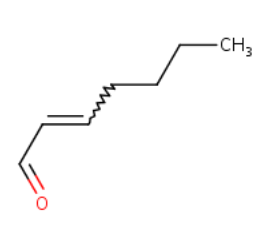 | C_7_H_12_O | 112.17 |
| 47 | Trans-2-Octenal | 5283324 | 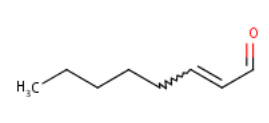 | C_8_H_14_O | 126.20 |
| 48 | Trans-2-Nonenal | 5283335 | 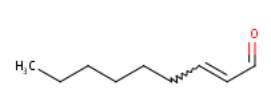 | C_9_H_16_O | 140.22 |
| 49 | Trans-2-Decenal | 5283345 | 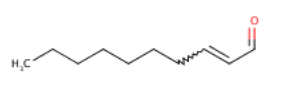 | C_10_H_18_O | 154.25 |
| 50 | Trans-2-Undecenal | 5283356 | 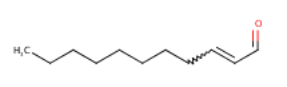 | C_11_H_20_O | 168.28 |
| 51 | Trans-2-Dodecenal | 5283361 | 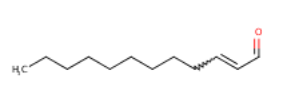 | C_12_H_22_O | 182.30 |
| 52 | Trans-2-Tridecenal | 5283363 | 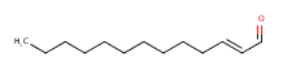 | C_13_H_24_O | 196.33 |
| 53 | Trans-3-Octen-2-one | 5363229 | 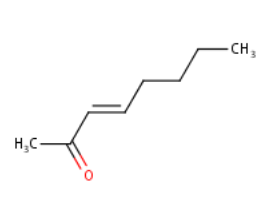 | C_8_H_14_O | 126.20 |
| 54 | Trans-3-Decen-2-one | 5363233 | 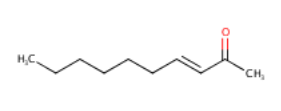 | C_10_H_18_O | 154.25 |
| 55 | Trans-3-Nonen-2-one | 5317045 | 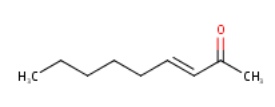 | C_9_H_16_O | 140.22 |
| 56 | 2-Octenoic acid | 5282714 | 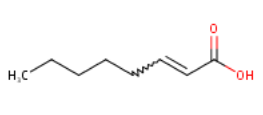 | C_8_H_14_O_2_ | 142.20 |
| 57 | Cis-3-None-1-ol | 5364631 | 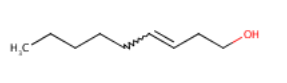 | C_9_H_18_O | 142.24 |
| 58 | Phenylethyl alcohol | 6054 | 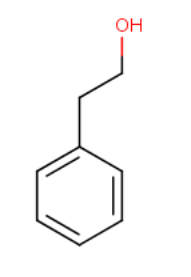 | C_8_H_10_O | 122.16 |
| 59 | Estragol | 8815 | 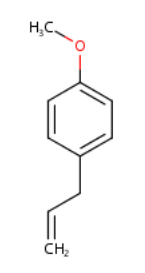 | C_10_H_12_O | 148.20 |
| 60 | p-Anisaldehyde | 31244 | 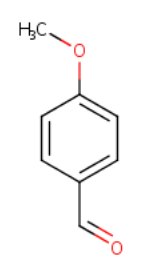 | C_8_H_8_O_2_ | 136.15 |
| 61 | 4-Hydroxy-2, 5-dimethyl-3(2H)-furanone | 11007913 | 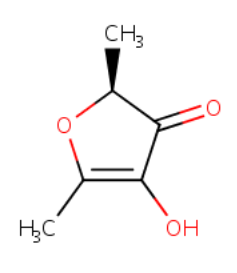 | C_6_H_8_O_3_ | 128.13 |
| 62 | 5, 8a-Di-1-propyl-octahydronaphthalen-1-(2H)-one | - | 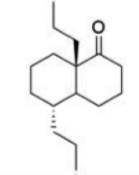 | - | - |
| 63 | Gnaphaliol 3-O-β-D-glucopyranoside | - | 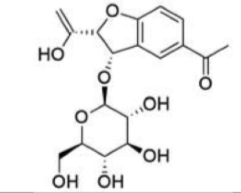 | - | - |
| 64 | Gnaphaliol 9-O-β-D-glucopyranoside | - | 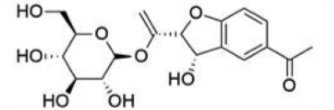 | - | - |
| 65 | Malvidin | 159287 | 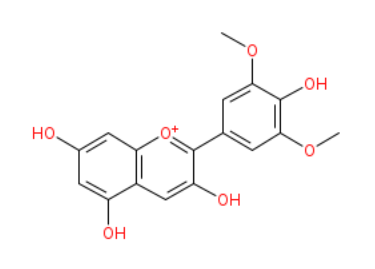 | C_17_H_15_O_7_ | 331.30 |
| 66 | Coumarin | 323 | 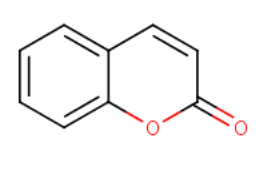 | C_9_H_6_O_2_ | 146.14 |
| 67 | Umbelliferone  (7-hydroxycoumarine) | 5281426 | 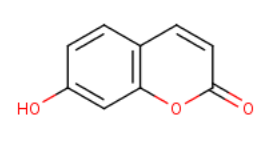 | C_9_H_6_O_3_ | 162.14 |
| 68 | Esculetin | 5281416 | 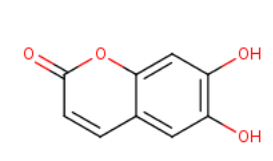 | C_9_H_6_O_4_ | 178.14 |
| 69 | Esculin | 5281417 | 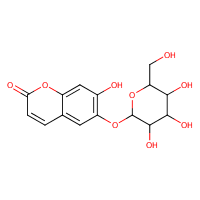 | C_15_H_16_O_9_ | 340.28 |
| 70 | Dimethyl-esculetin | 8417 | 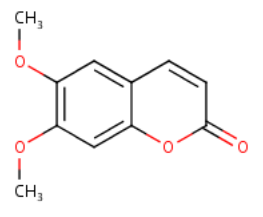 | C_11_H_10_O_4_ | 206.19 |
| 71 | Psoralen | 6199 | 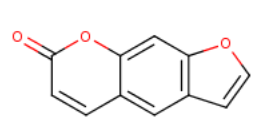 | C_11_H_6_O_3_ | 186.16 |
| 72 | Nodakenetin | 26305 | 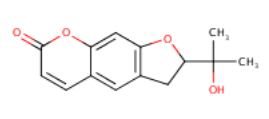 | C_14_H_14_O_4_ | 246.26 |
| 73 | Coladonin | 7067264 | 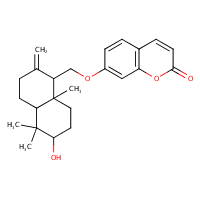 | C_24_H_30_O_4_ | 382.5 |
| 74 | Dihydroxy-bergamottin | 12082365 | 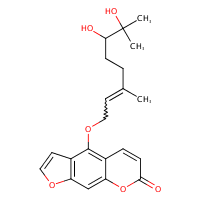 | C_21_H_24_O_6_ | 372.4 |
| 75 | Bergamottin | 5471349 | 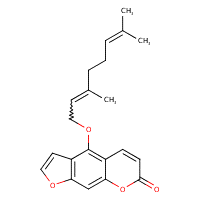 | C_21_H_22_O_4_ | 338.4 |
| 76 | Imperatorin | 10212 | 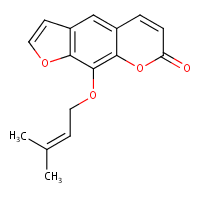 | C_16_H_14_O_4_ | 270.28 |
| 77 | Chalcone | 637760 | 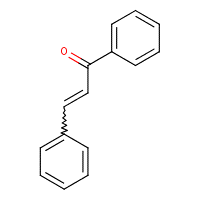 | C_15_H_12_O | 208.25 |
| 78 | 2’, 4’- Dihydroxychalcone | 5376979 | 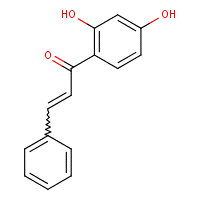 | C_15_H_12_O_3_ | 240.25 |
| 79 | 2, 2’, 4’- Trihydroxychalcone | 5811533 | 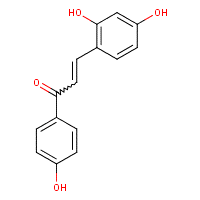 | C_15_H_12_O_4_ | 256.25 |
| 80 | 2’, 4’-Dihydroxy-2-mehoxychalcone | - | 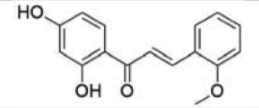 | - | - |
| 81 | *Trans*-Benzylidene acetophenone | 637760 | 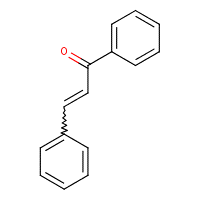 | C_15_H_12_O | 208.25 |
| 82 | Phloretin | 4788 | 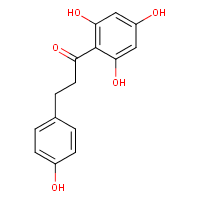 | C_15_H_14_O_5_ | 274.27 |
| 83 | Phloridzin | 6072 | 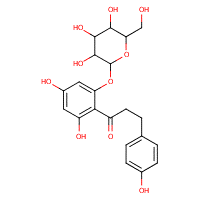 | C_21_H_24_O_10_ | 436.4 |
| 84 | 2’, 3’, 5- Trihydroxy-4’, 6’, 3-trimethoxychalcone | - | 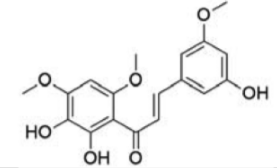 | - | - |
| 85 | 2’, 3’-Dihydroxy-4’, 6’-dimethoxychalcone | - | 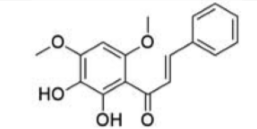 | - | - |
| 86 | 2’, 4’, 4-Trihydroxy-3, 6’- dimethoxychalcone | - | 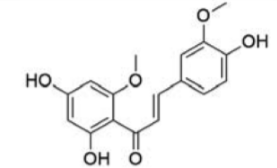 | - | - |
| 87 | Xanthohumol | 639665 | 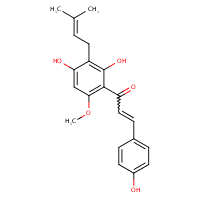 | C_21_H_22_O_5_ | 354.4 |
| 88 | Licochalcone A | 5318998 | 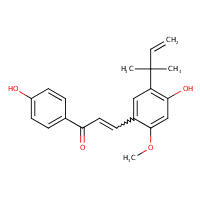 | C_21_H_22_O_4_ | 338.4 |
| 89 | Licochalcone E | 46209991 | 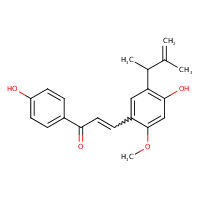 | C_21_H_22_O_4_ | 338.40 |
| 90 | Catechin | 9064 | 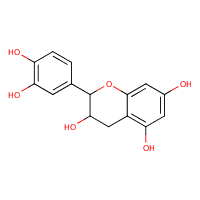 | C_15_H_14_O_6_ | 290.27 |
| 91 | Epicatechin | 72276 | 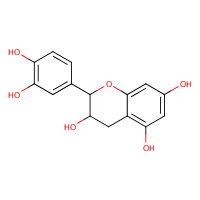 | C_15_H_14_O_6_ | 290.27 |
| 92 | Epicatechin gallate | 107905 | 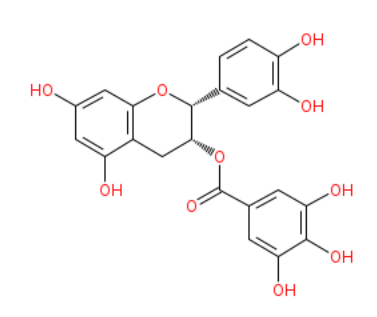 | C_22_H_18_O_10_ | 442.37 |
| 93 | Epigallocatechin | 72277 | 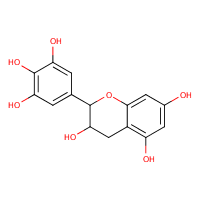 | C_15_H_14_O_7_ | 306.27 |
| 94 | Epigallocatechin gallate | 65064 | 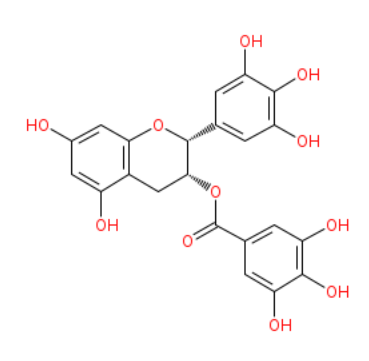 | C_22_H_18_O_11_ | 458.37 |
| 95 | (-)-(2S)-7, 5’-Dihydroxy- 5, 3’-dimethoxyflavanone | - | 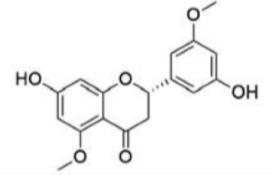 | - | - |
| 96 | Glabranine | 124049 | 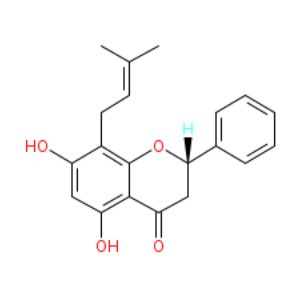 | C_20_H_20_O_4_ | 324.37 |
| 97 | Naringenin | 932 | 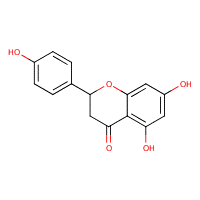 | C_15_H_12_O_5_ | 272.25 |
| 98 | Naringin | 442428 | 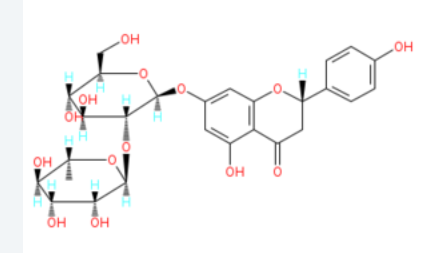 | C_27_H_32_O_14_ | 580.53 |
| 99 | 8-Prenylnaringnin | 480764 | 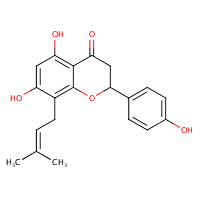 | C_20_H_20_O_5_ | 340.4 |
| 100 | Isosakuranetin | 160481 | 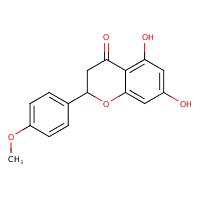 | C_16_H_14_O_5_ | 286.28 |
| 101 | Farrerol | 91144 |  | C_17_H_16_O_5_ | 300.3 |
| 102 | Cyrtominetin | 125309 |  | C_17_H_16_O_6_ | 316.3 |
| 103 | Liquiritigenin | 114829 |  | C_15_H_12_O_4_ | 256.25 |
| 104 | Pinocembrin | 68071 |  | C_4_H_12_O_4_ | 256.25 |
| 105 | Eriodictyol | 440735 |  | C_15_H_12_O_6_ | 288.25 |
| 106 | Amoradicin | 11178127 |  | C_26_H_30_O_6_ | 438.51 |
| 107 | Amorisin | 16727161 |  | C_30_H_36_O_6_ | 492.60 |
| 108 | Isoamoritin | 42607986 |  | C_31_H_38_O_6_ | 506.63 |
| 109 | Amoricin | 14187653 |  | C_31_H_36_O_6_ | 504.61 |
| 110 | Kurarinol | 44563198 |  | C_26_H_32_O_7_ | 456.5 |
| 111 | Alopecurone H | - |  | - | - |
| 112 | Alopecurone I | - |  | - | - |
| 113 | Alopecurone J | - |  | - | - |
| 114 | Alopecurone D | 101915836 |  | C_40_H_40_O_9_ | 664.74 |
| 115 | Alopecurone A | 10032196 |  | C_39_H_38_O_9_ | 650.71 |
| 116 | Hesperidin | 10621 |  | C_28_H_34_O_15_ | 610.56 |
| 117 | Neohesperidin | 442439 |  | C_28_H_34_O_15_ | 610.56 |
| 118 | Neoriocitrin | - |  | - | - |
| 119 | Taxifolin | 439533 |  | C_15_H_12_O_7_ | 304.25 |
| 120 | Flavone | 10680 |  | C_15_H_10_O_2_ | 222.24 |
| 121 | 6-Hydroxyflavone | 72279 |  | C_15_H_10_O_3_ | 238.24 |
| 122 | 6-Aminoflavone | 16217320 |  | C_15_H_11_NO_2_ | 237.25 |
| 123 | Apigenin | 5280443 |  | C_15_H_10_O_5_ | 270.24 |
| 124 | Chrysin | 5281607 |  | C_15_H_10_O_4_ | 254.24 |
| 125 | Luteolin | 5280445 |  | C_15_H_10_O_6_ | 286.24 |
| 126 | Baicalein | 5281605 |  | C_15_H_10_O_5_ | 270.24 |
| 127 | Baicalin | 64982 |  | C_21_H_18_O_11_ | 446.36 |
| 128 | Oroxylin A | 5320315 |  | C_16_H_12_O_5_ | 284.26 |
| 129 | Oroxylin A 7-*O*-glucoronide | 14655552 |  | C_22_H_20_O_11_ | 460.4 |
| 130 | Oroxylin B | - |  | - | - |
| 131 | 3’, 4’, 5-Trihydroxy-6, 7-dimethoxy-flavone | - |  | - | - |
| 132 | 5, 6, 7, 3, 4’-Pentahydroxy-flavone | - |  | - | - |
| 133 | Heptamethoxy-flavone | 150893 |  | C22H24O9 | 432.42 |
| 134 | Nobiletin | 72344 |  | C21H22O8 | 402.39 |
| 135 | Sinesetin | 145659 |  | C_20_H_20_O_7_ | 372.37 |
| 136 | Wogonoside | 3084961 |  | C_22_H_20_O_11_ | 460.39 |
| 137 | Artocarpin | 5458461 |  | C_26_H_28_O_6_ | 436.50 |
| 138 | Scutellarin | 185617 |  | C_21_H_18_O_12_ | 462.36 |
| 139 | Icariin | 5318997 |  | C_33_H_40_O_15_ | 676.66 |
| 140 | Isovitexin | 162350 |  | C_21_H_20_O_10_ | 432.38 |
| 141 | Vitexin | 5280441 |  | C_21_H_20_O_10_ | 432.38 |
| 142 | Isoorientin | 114776 |  | C_21_H_20_O_11_ | 448.38 |
| 143 | Orientin | 5281675 |  | C_21_H_20_O_11_ | 448.38 |
| 144 | Quercetin | 5280343 |  | C_15_H_10_O_7_ | 302.23 |
| 145 | Quercetin-3O-arabinosid | - |  | - | - |
| 146 | Quercitrin | 5280459 |  | C_21_H_20_O_11_ | 448.38 |
| 147 | Myricetin | 5281672 |  | C_15_H_10_O_8_ | 318.24 |
| 148 | Morin | 5281670 |  | C_15_H_10_O_7_ | 302.23 |
| 149 | Fisetin | 5281614 |  | C_15_H_10_O_6_ | 286.24 |
| 150 | Kaempferol | 5280863 |  | C_15_H_10_O_6_ | 286.24 |
| 151 | Kaempferol-3-rutinoside | 5318767 |  | C_27_H_30_O_15_ | 594.52 |
| 152 | Isorhamnetin-3-*O-B*-D-rutinoside | - |  | - | - |
| 153 | Rutin | 5280805 |  | C_27_H_30_O_16_ | 610.52 |
| 154 | Silibinin | 31553 |  | C_25_H_22_O_10_ | 482.44 |
| 155 | Daidzein | 5281708 |  | C_15_H_10_O_4_ | 254.24 |
| 156 | Genistein | 5280961 |  | C_15_H_10_O_5_ | 270.24 |
| 157 | 8-y,y-Dimethyl-allylwighteone | - |  | - | - |
| 158 | Flemingsin | 15719494 |  | C_26_H_28_O_6_ | 436.50 |
| 159 | 6, 8- Diprenylorobol | 21148065 |  | C_25_H_26_O_6_ | 422.5 |
| 160 | Auriculasin | 5358846 |  | C_25_H_24_O_6_ | 420.45 |
| 161 | Flemiphilippinin A | 10074228 |  | C30H32O6 | 488.57 |
| 162 | Flemiphilippinin E | - |  | - | - |
| 163 | Osajin | 95168 |  | C_25_H_24_O_5_ | 404.46 |
| 164 | 5, 7, 30-Tetrahydroxy-20, 50-di(3-mthylbut-2-enyl)isoflavone | - |  | - | - |
| 165 | 5, 7, 30-Trihydroxy-20-(3-methylbut-2-enyl)-40, 50-(3, 3-dimethylpyrano) isoflavone | - |  | - | - |
| 166 | Puerarin | 5281807 |  | C_21_H_20_O_9_ | 416.4 |
| 167 | Demethylmedicarpin | 162933 |  | C_15_H_12_O_4_ | 256.25 |
| 168 | Neorautenol | 11500744 |  | C_20_H_18_O_4_ | 322.35 |
| 169 | Isoneorautenol | 73649 |  | C_20_H_18_O_4_ | 322.35 |
| 170 | Phaseollin | 91572 |  | C_20_H_18_O_4_ | 322.4 |
| 171 | Eryvarin D | 15546808 |  | C_21_H_20_O_4_ | 336.38 |
| 172 | Erythribyssin O | 46861837 |  | C_21_H_18_O_5_ | 3509.36 |
| 173 | Calopocarpin | 11709595 |  | C_20_H_20_O_4_ | 324.37 |
| 174 | Erythribyssin L | 46887764 |  | C_25_H_28_O_5_ | 408.49 |
| 175 | Erysubin D | 12051846 |  | C_25_H_28_O_5_ | 408.49 |
| 176 | Erysubin E | 637080 |  | C_25_H_26_O_5_ | 406.47 |
| 177 | Erythribyssin D | 46887822 |  | C_20_H_20_O_5_ | 340.37 |
| 178 | Erythribyssin M | 46887823 |  | C_20_H_20_O_5_ | 340.4 |
| 179 | Cristacarpin | 126540 |  | C_21_H_22_O_5_ | 354.40 |
| 180 | Sophorapterocarpan A | 14017299 |  | C_20_H_20_O_4_ | 324.27 |
| 181 | Erystagallin A | 10410005 |  | C_26_H_30_O_5_ | 422.51 |
| 182 | Bicolosin A | 56668791 |  | C_27_H_32_O_5_ | 436.54 |
| 183 | Bicolosin B | 56665359 |  | C_31_H_38_O_4_ | 474.63 |
| 184 | Bicolosin C | 56682289 |  | C_26_H_30_O_5_ | 422.51 |
| 185 | Erythrabyssin II | 10408212 |  | C_25_H_28_O_4_ | 392.5 |
| 186 | Lespebuergine G4 | 56657670 |  | C_26_H_30_O_4_ | 406.51 |
| 187 | 1-Methoxy-erythrabyssin II | - |  | - | - |
| 188 | Amorphigenin | 92207 |  | C_23_H_22_O_7_ | 410.42 |
| 189 | Dalbinol | 44257412 |  | C_23_H_22_O_8_ | 426.4 |
| 190 | 6-Ketodehydro-amorphigenin | - |  | - | - |
| 191 | Macelignan | 10404245 |  | C_20_H_24_O_4_ | 328.40 |
| 192 | Magnolol | 72300 |  | C_18_H_18_O_2_ | 266.33 |
| 193 | Medioresinol | 181681 |  | C_21_H_24_O_7_ | 388.41 |
| 194 | (7S, 8S)-Dihydro-dehydrodiconiferyl alcohol | - |  | - | - |
| 195 | (7S, 8S)-Dihydro- dehydrodiconiferyl alcohol 9’-O-β-D-glucopyranoside | - |  | - | - |
| 196 | Gallic acid | 370 |  | C_7_H_6_O_5_ | 170.12 |
| 197 | Methyl gallate | 7428 |  | C_8_H_8_O_5_ | 184.15 |
| 198 | Sallicylic acid | 338 |  | C_7_H_6_O_3_ | 138.12 |
| 199 | p-Hydroxybenzoic acid | 135 |  | C_7_H_6_O_3_ | 138.12 |
| 200 | Acetyl salicylic acid | 2244 |  | C_9_H_8_O_4_ | 180.16 |
| 201 | Methyl salicylate | 4133 |  | C_8_H_8_O_3_ | 152.15 |
| 202 | Salicylamide | 5147 |  | C_7_H_7_NO_2_ | 137.14 |
| 203 | Benzoic acid | 243 |  | C_7_H_6_O_2_ | 122.12 |
| 204 | Protocatechuic acid | 72 |  | C_7_H_6_O_4_ | 154.12 |
| 205 | Vanillic acid | 8468 |  | C_8_H_8_O_4_ | 168.15 |
| 206 | Ginkgolic acid C 15:1 | 5281858 |  | C_22_H_34_O_3_ | 346.50 |
| 207 | Ginkgolic acid C 17:1 | 5469634 |  | C_24_H_38_O_3_ | 374.56 |
| 208 | Malabaricone C | 100313 |  | C_21_H_26_O_5_ | 358.43 |
| 209 | 4-Hydroxytyrosol | - |  | - | - |
| 210 | Salidroside | 159278 |  | C_14_H_20_O_7_ | 300.3 |
| 211 | Desmethylyangonnine-4’-*O*-[6’’-*O*-(3-hydroxy-3-methylglutaryl)]-B-D-glucopyranoside | - |  | - | - |
| 212 | Desmethylyangonnine-4’-*O*-[6”-O-malonyl)-B-D-glucopyranoside | - |  | - | - |
| 213 | Desmethylyangonnine-4’-O-B-D-glucopyranoside | - |  | - | - |
| 214 | Maltol 3-O-(4’-O-p-coumaroyl-6’-O-(3-hydroxy-3-methylglutaroyl))-B- glucopyranoside | - |  | - | - |
| 215 | Maltol-3-O-(4’-O-cis-p-coumaroyl-6’-O-(3-hydroxy-3-methylglutaroyl))-B-glucopyranoside | - |  | - | - |
| 216 | Oleuropein glucoside | - |  | - | - |
| 217 | p-Coumaric acid | 637542 |  | C_9_H_8_O_3_ | 164.16 |
| 218 | Caffeic acid | 689043 |  | C_9_H_8_O_4_ | 180.16 |
| 219 | Ferulic acid | 445858 |  | C_10_H_10_O_4_ | 194.18 |
| 220 | Cinnamic acid | 444539 |  | C_9_H_8_O_2_ | 148.16 |
| 221 | Cinnamaldehyde | 637511 |  | C_9_H_8_O | 132.16 |
| 222 | 4-Methoxy cinnamaldehyde | 641294 |  | C_10_H_10_O_2_ | 162.19 |
| 223 | 2-Methoxi cinnamaldehyde | 641298 |  | C_10_H_10_O_2_ | 162.19 |
| 224 | 4-Dimethylamino-cinnamaldehyde | 92224 |  | C_11_H_13_NO | 175.23 |
| 225 | 4-Phenyl-2-butanone | 17355 |  | C_10_H_12_O | 148.20 |
| 226 | Eugenol | 3314 |  | C_10_H_12_O_2_ | 164.20 |
| 227 | Isoeugenol | 853433 |  | C_10_H_12_O_2_ | 164.20 |
| 228 | Methyl eugenol | 7127 |  | C_11_H_14_O_2_ | 178.23 |
| 229 | Eugenyl acetate | 7136 |  | C_12_H_14_O_3_ | 206.24 |
| 230 | Chlorogenic acid | 1794427 |  | C_16_H_18_O_9_ | 354.31 |
| 231 | Rosmarinic acid | 5281792 |  | C_18_H_16_O_8_ | 360.3 |
| 232 | 3,5-Dicaffeoylquinic acid | 6474310 |  | C_25_H_24_O_12_ | 516.4 |
| 233 | Nordihydroguaiaretic acid | 4534 |  | C_18_H_22_O_4_ | 302.4 |
| 234 | 6-Gingerol | 442793 |  | C_17_H_26_O_4_ | 294.4 |
| 235 | Zingerone | 31211 |  | C_11_H_14_O_3_ | 194.23 |
| 236 | 6-Shogaol | 5281794 |  | C_17_H_24_O_3_ | 276.37 |
| 237 | Curcumin | 969516 |  | C_21_H_20_O_6_ | 268.38 |
| 238 | Demethoxy-curcumin | 5469424 |  | C_20_H_18_O_5_ | 338.4 |
| 239 | Bisdemethoxy-curcumin | 5315472 |  | C_19_H_16_O_4_ | 308.3 |
| 240 | Katsumadain A | 9982435 |  | C_32_H_28_O_4_ | 476.56 |
| 241 | Emodin | 3220 |  | C_15_H_10_O_5_ | 270.24 |
| 242 | Hypericin | 3663 |  | C_30_H_16_O_8_ | 504.44 |
| 243 | Quinone | 4650 |  | C_6_H_4_O_2_ | 108.09 |
| 244 | Chrysophanol | 10208 |  | C_15_H_10_O_4_ | 254.24 |
| 245 | Thymoquinone | 10281 |  | C_10_H_12_O_2_ | 164.20 |
| 246 | 10’(Z), 13’(E)-Heptadecadienylhydroquinone | 10991557 |  | C_23_H_36_O_2_ | 344.53 |
| 247 | Shikonin | 479503 |  | C_16_H_16_O_5_ | 288.29 |
| 248 | Resorcinol | 5054 |  | C_6_H_6_O_2_ | 110.11 |
| 249 | 3-Geranyl-1-(2-methylpropanoylphloroglucinol) | 10426888 |  | C_20_H_28_O_4_ | 332.4 |
| 250 | 3-Geranyl-1-(2-methylbutanoyl) phloroglucinol | 14282627 |  | C_21_H_30_O_4_ | 346.46 |
| 251 | 2-Geranyloxy-1-(2-methypropanoyl)phloroglucinol | - |  | - | - |
| 252 | Panduratin A | 6483648 |  | C_26_H_30_O_4_ | 406.5 |
| 253 | Rhodomyrtone | 12050020 |  | C_26_H_34_O_6_ | 442.5 |
| 254 | 7-Epiclusianone | 5471610 |  | C_33_H_42_O_4_ | 502.68 |
| 255 | Hyperforin | 441298 |  | C_35_H_52_O_4_ | 536.78 |
| 256 | Dihydroxybenzofurane | - |  | - | - |
| 257 | *Cis*-stilbene | 5356785 |  | C_14_H_12_ | 180.25 |
| 258 | *Trans*-stilbene | 638088 |  | C_14_H_12_ | 180.12 |
| 259 | Dicinnamyl | 5376733 |  | C_18_H_16_ | 232.32 |
| 260 | Resveratrol | 445154 |  | C_14_H_12_O_3_ | 228.24 |
| 261 | Oxyresverartrol | 5281717 |  | C_14_H_12_O_4_ | 244.24 |
| 262 | ɛ-Viniferin | 11236373 |  | C_28_H_22_O_6_ | 454.47 |
| 263 | Suffruticosol A | 101203677 |  | C_47_H_40_O_9_ | 748.82 |
| 264 | Suffruticosol B | 101203678 |  | C_47_H_40_O_9_ | 748.82 |
| 265 | Vitisin A | 16131430 |  | C_56_H_42_O_12_ | 906.93 |
| 266 | Vitisin B | 74947464 |  | C_25_H_25_O_12_ | 517.458 |
| 267 | *Trans*-Gnetin H | 9852931 |  | C_42_H_32_O_9_ | 680.70 |
| 268 | Ellagic acid | 5281855 |  | C_14_H_6_O_8_ | 302.19 |
| 269 | 3-*O*-Methyl ellagic acid | 78384860 |  | C_20_H_16_O_12_ | 448.33 |
| 270 | 2,5-Di-O-galloyl-D-hamamelose  (Hamemelitannin) | 44584241 |  | C_20_H_20_O_14_ | 484.36 |
| 271 | 1,2,3,4,6-Penta-O-galloy-B-D-glucopyranose | 65238 |  | C_41_H_32_O_26_ | 940.67 |
| 272 | Tannic acid | 16129778 |  | C_76_H_52_O_46_ | 1701.2 |
| 273 | Punicalagin | 44584733 |  | C_48_H_28_O_30_ | 1084.72 |
| 274 | Proanthocyanidin A2 | 124025 |  | C_30_H_24_O_12_ | 576.50 |
| 275 | Proanthocyanidin B2 | 122738 |  | C_30_H_26_O_12_ | 578.52 |
| 276 | Proanthocyanidin B3 | 146798 |  | C_30_H_26_O_12_ | 578.52 |
| 277 | Proanthocyanidin C1 | 169853 |  | C_45_H_38_O_18_ | 866.77 |
| 278 | Proanthocyanidin C2 | 11182062 |  | C_45_H_38_O_18_ | 866.77 |
| 279 | B-type linked proanthocyanidins | - |  | - | - |
| 280 | B-type linked prothocyanidins | - |  | - | - |
| 281 | A and B-type linked proanthocyanidins | - |  | - | - |
| 282 | A-type proanthocyanidins (from trimers to oligomers with high polymerization degree) | - |  | - | - |
| 283 | Mangostanta-xanthone I | - |  | - | - |
| 284 | α-Mangostin | 5281650 |  | C_24_H_26_O_6_ | 410.46 |
| 285 | Mangiferin | 5281647 |  | C_19_H_18_O_11_ | 422.34 |
| 286 | β-Sitosterol-3-O-glucopyranoside | 296119 |  | C_35_H_60_O_6_ | 576.85 |
| 287 | α-Phellandrene | 443160 |  | C_10_H_16_ | 136.23 |
| 288 | *p*-Cymene | 7463 |  | C_10_H_14_ | 134.22 |
| 289 | Thymol | 6989 |  | C_10_H_14_O | 150.22 |
| 290 | Carvacrol | 10364 |  | C_10_H_14_O | 150.22 |
| 291 | α-Terpineol | 442501 |  | C_10_H_18_O | 154.25 |
| 292 | Thujone | 261491 |  | C_10_H_16_O | 152.23 |
| 293 | Citral | 638011 |  | C_10_H_16_O | 152.23 |
| 294 | Citronellol | 8842 |  | C_10_H_20_O | 156.27 |
| 295 | Citronellal | 7794 |  | C_10_H_18_O | 154.25 |
| 296 | Geraniol | 637566 |  | C_10_H_18_O | 154.25 |
| 297 | (-)-Carvone | 439570 |  | C_10_H_14_O | 150.22 |
| 298 | Pulegone | 442495 |  | C_10_H_16_O | 152.23 |
| 299 | Menthone | 26447 |  | C_10_H_18_O | 154.25 |
| 300 | Menthol | 1254 |  | C_10_H_20_O | 156.27 |
| 301 | Terpin | 6651 |  | C_10_H_20_O_2_ | 172.26 |
| 302 | Limonene | 22311 |  | C_10_H_16_ | 136.23 |
| 303 | Terpinene-4-ol | 11230 |  | C_10_H_18_O | 154.25 |
| 304 | 1, 8-Cineol | 2758 |  | C_10_H_18_O | 154.25 |
| 305 | Linalool | 6549 |  | C_10_H_18_O | 154.25 |
| 306 | α-Pinene | 440968 |  | C_10_H_16_ | 136.23 |
| 307 | Camphene | 6616 |  | C_10_H_16_ | 136.23 |
| 308 | Camphor | 2537 |  | C_10_H_16_O | 152.23 |
| 309 | (-) Borneol | 1201518 |  | C_10_H_18_O | 154.25 |
| 310 | Phytol | 5280435 |  | C_20_H_40_O | 296.53 |
| 311 | Geranyllinalool | 5365872 |  | C_20_H_34_O | 290.48 |
| 312 | Dehydroabieic acid | 94391 |  | C_20_H_28_O_2_ | 300.44 |
| 313 | Kaurenoic acid | 73062 |  | C_20_H_30_O_2_ | 302.45 |
| 314 | *Ent*-trachyloban-19-oic acid | 23601154 |  | C_20_H_30_O_2_ | 302.45 |
| 315 | Casbane diterpene (1, 4-dihydroxy-2E, 6E, 12E-trien-5-one-casbane | - |  | - | - |
| 316 | Andensin | 14707350 |  | C_20_H_32_O_2_ | 304.5 |
| 317 | Xanthorrhizol | 93135 |  | C_15_H_22_O | 218.33 |
| 318 | Farnesol | 445070 |  | C_15_H_26_O | 222.37 |
| 319 | Farnesyl acetate | 94403 |  | C_17_H_28_O_2_ | 264.40 |
| 320 | Nerolidol | 5284507 |  | C_15_H_26_O | 222.37 |
| 321 | Nerol | 643820 |  | C_10_H_18_O | 154.25 |
| 322 | Valencene | 9855795 |  | C_15_H_24_ | 204.35 |
| 323 | α-Cyperone | 12303268 |  | C_15_H_22_O | 218.33 |
| 324 | Isoalantolactone | 73285 |  | C_15_H_20_O_2_ | 232.32 |
| 325 | Patchouli alcohol | 10955174 |  | C_15_H_26_O | 222.37 |
| 326 | *Ent*-spathulenol | 13854255 |  | C_15_H_24_O | 220.35 |
| 327 | *Ent*-4β, 10α-dihydroxy-aromadendrane | - |  | - | - |
| 328 | Viridiflorol | 11996452 |  | C_15_H_26_O | 222.37 |
| 329 | 1-Oxo-3, 10-epoxy-5-hydroxy-8-metacryloyloxy-germacra-2, 4(15), 11(13)-trien-6, 12-olide | - |  | - | - |
| 330 | 1-Oxo-3, 10-epoxy-8-methacryloyloxy-15-hydroxygermacra-2, 4, 11(13)-trien-6, 12-olide | - |  | - | - |
| 331 | 1-Oxo-3, 10-epoxy-8-epoxymethacryloyloxy-15-hydroxygermacra-2, 4, 11(13)-trien-6, 12-olide | - |  | - | - |
| 332 | 1-Oxo-3, 10-epoxy-5-hydroxy-8-angeloyloxy-germica-2, 4(15), 11(13)-trien-6, 12-olide | - |  | - | - |
| 333 | 1-Oxo-3, 10-epoxy-8-angeloyloxy-15-hydroxygermacra-2,4,11(13)-trien-6,12-olide | - |  | - | - |
| 334 | 1-Oxo-3, 10-epoxy-5-hydroxy-8-tigloyloxy-germacra-2,4(15), 11(13)-trien-6, 12-olide | - |  | - | - |
| 335 | 5-Epidilatanolide A | 75717755 |  | - | - |
| 336 | 5-Epidilatanolide B | 79098238 |  | - | - |
| 337 | Lecocarpinolide B | 101630355 |  | C_22_H_26_O_8_ | 418.44 |
| 338 | 9r-Acetyloxy-8β-angeloyloxy-14-hydroxy-acanthospermolide | - |  | - | - |
| 339 | 9r-Acetyloxy-14, 15-dihydroxy-8β-(2 –methoxylbutanoyloxy)-acanthospermolide | - |  | - | - |
| 340 | 9r-Acetyloxy- 14, 15-dihydroxy-8β-angeloyloxy-Acanthospermolide | - |  | - | - |
| 341 | 19-Hydroxy-15-desoxy-orientalide | - |  | - | - |
| 342 | 15-Acetoyloxy-8β-isobutanoyloxy-14-oxo-(4Z)-acanthospermolide | - |  | - | - |
| 343 | 15-Acetyloxy-8β-angeloyloxy-14-oxo-(4Z)-acanthospermolide | - |  | - | - |
| 344 | Isolimonic acid | 25022666 |  | C_26_H_32_O_9_ | 488.53 |
| 345 | Ichangin | 441801 |  | C_26_H_32_O_9_ | 488.53 |
| 346 | Isoobacunoic acid | 20055680 |  | C26H32O8 | 472.53 |
| 347 | Isoobacunoic acid glucoside | - |  | - | - |
| 348 | Deacetyl nomilinic acid glucoside | - |  | - | - |
| 349 | Limonin | 179651 |  | C_26_H_30_O_8_ | 470.51 |
| 350 | Obacunone | 119041 |  | C_26_H_30_O_7_ | 454.51 |
| 351 | Nomilin | 72320 |  | C_28_H_34_O_9_ | 514.56 |
| 352 | Deacetylnomilin | 13857953 |  | C_26_H_32_O_8_ | 472.53 |
| 353 | Limonin 17-β-D-glucopyranoside | 423545083 |  | - | - |
| 354 | 3 β, 6 β, 16 β- Trihydroxylup-20(29)-ene | - |  | - | - |
| 355 | Betulinic acid | 64971 |  | C_30_H_48_O_3_ | 456.70 |
| 356 | Asiatic acid | 119034 |  | C_30_H_48_O_5_ | 488.70 |
| 357 | Corosolic acid | 6918774 |  | C_30_H_48_O_4_ | 472.70 |
| 358 | Ursolic acid | 64945 |  | C_30_H_48_O_3_ | 456.70 |
| 359 | 3β-*O-Cis-p*-coumaroyl-20β-hydroxy-12-ursen-28-oic acid | - |  | - | - |
| 360 | 3β-*O-Trans-p-*coumaroyl-2α hydroxyl-12-ursen-28-oic-acid | - |  | - | - |
| 361 | 3β-*O-Cis-p-* coumaroyl-2α hydroxyl-12-ursen-28-oic-acid | - |  | - | - |
| 362 | 3β-*O-Trans-*feruloyl-2αhydroxy-12-ursen-28-28-oic acid | - |  | - | - |
| 363 | Celastrol | 122724 |  | C_29_H_38_O_4_ | 450.61 |
| 364 | Glycyrrhetinic acid | 10114 |  | C_30_H_46_O_4_ | 470.68 |
| 365 | Taraxerol | 92097 |  | C_30_H_50_O | 426.72 |
| 366 | Oleanolic acid | 10494 |  | C_30_H_48_O_3_ | 456.70 |
| 367 | Shoreic acid | 12315515 |  | C_30_H_50_O_4_ | 474.72 |
| 368 | Eichlerialactone | 76313961 |  | C_27_H_42_O_4_ | 460.32 |
| 369 | Cabraleone | 21625900 |  | C_30_H_50_O_3_ | 458.72 |
| 370 | Cabraleadiol | 21625899 |  | C_30_H_52_O_3_ | 460.73 |
| 371 | Atractylenolide III | 155948 |  | \| [C_15_H_20_O_3_](https://pubchem.ncbi.nlm.nih.gov#query=C15H20O3) | 248.32 |
| 372 | Epiestriol | 68929 |  | C_18_H_24_O_3_ | 288.4 |
| 373 | Phycoerythrobilin | 20057005 |  | [C_33_H_38_N_4_O_6_](https://pubchem.ncbi.nlm.nih.gov#query=C33H38N4O6) | 586.7 |
| 374 | Phycocyanobilin | 365902 |  | [C_33_H_38_N_4_O_6_](https://pubchem.ncbi.nlm.nih.gov#query=C33H38N4O6) | 586.7 |
| 375 | Dieckol | 3008868 |  | [C_36_H_22_O_18_](https://pubchem.ncbi.nlm.nih.gov#query=C36H22O18) | 742.5 |
| 376 | Phycourobilin | 5289229 |  | [C_33_H_42_N_4_O_6_](https://pubchem.ncbi.nlm.nih.gov#query=C33H42N4O6) | 590.7 |
| 377 | Folic acid | 135398658 |  | [C_19_H_19_N_7_O_6_](https://pubchem.ncbi.nlm.nih.gov#query=C19H19N7O6) | 441.4 |
| 378 | β-carotene | 5280489 |  | [C_40_H_56_](https://pubchem.ncbi.nlm.nih.gov#query=C40H56) | 536.9 |
| 370 | Astaxanthin | 5281224 |  | [C_40_H_52_O_4_](https://pubchem.ncbi.nlm.nih.gov#query=C40H52O4) | 596.8 |
| 380 | Aloe-emodin | 10207 |  | [C_15_H_10_O_5_](https://pubchem.ncbi.nlm.nih.gov#query=C15H10O5) | 270.24 |
| 381 | Anthrarufin | 8328 |  | [C_14_H_8_O_4_](https://pubchem.ncbi.nlm.nih.gov#query=C14H8O4) | 240.21 |
| 382 | Alizarin | 6293 |  | [C_14_H_8_O_4_](https://pubchem.ncbi.nlm.nih.gov#query=C14H8O4) | 240.21 |
| 383 | Dantron | 2950 |  | [C_14_H_8_O_4_](https://pubchem.ncbi.nlm.nih.gov#query=C14H8O4) | 240.21 |
| 384 | 3-β-Hydroxy-nordammaran-20-one | - |  | - | - |

**Table S3.** The obtained models for OmpA (**A**) and OmpW (**B**) by GalaxyRefine server.

**A**

**B**

**Table S4.** Physicochemical properties and adequate absorption, distribution, metabolism, excretion and tolerable toxicity (ADMET) evaluation of the successfully filtered out potential drug molecules.

| Compound name | PubChem ID | Mw | nRot | HBD | HBA | TPSA (Å2) | logP | MR | GI absorption | Log Papp | Log BB | Log PS | LD50 (mol/kg) | Total clearance |
| --- | --- | --- | --- | --- | --- | --- | --- | --- | --- | --- | --- | --- | --- | --- |
| Indole-3-acetamide | 397 | 174.2 | 2 | 2 | 1 | 58.88 | 1.1 | 50.98 | High | 1.14 | -0.29 | -2.33 | 2.13 | 0.45 |
| Atractylenolide III | 155948 | 248.322 | 0 | 1 | 3 | 107.521 | 2.7 | 69.15 | High | 1.314 | 0.603 | -2.381 | 1.826 | 0.995 |
| Epiestriol | 68929 | 288.4 | 0 | 3 | 3 | 125.176 | 2.58 | 82.19 | High | 1.262 | -0.218 | -2.095 | 2.649 | 0.789 |
| Aloe-emodin | 10207 | 270.24 | 1 | 3 | 5 | 113.283 | 1.36 | 69.92 | High | -0.217 | -0.729 | -2.466 | 2.329 | 0.008 |
| Anthrarufin | 8328 | 240.21 | 0 | 2 | 4 | 102.124 | 1.87 | 63.8 | High | 1.317 | 0.202 | -2.164 | 2.233 | 0.004 |
| Alizarin | 6293 | 240.21 | 0 | 2 | 4 | 102.124 | 1.87 | 63.8 | High | 1.075 | 0.184 | -2.157 | 2.255 | 0.046 |
| Dantron | 2950 | 240.21 | 0 | 2 | 4 | 102.124 | 1.87 | 63.8 | High | 1.058 | 0.202 | -2.164 | 2.233 | 0.005 |
| Oxyresveratrol | 5281717 | 244.24 | 2 | 4 | 4 | 103.706 | 2.67 | 69.9 | High | 0.831 | -0.899 | -2.304 | 2.354 | 0.195 |
| Psoralen | 6199 | 186.16 | 0 | 0 | 3 | 43.35 | 2.12 | 52.26 | High | 1.29 | 0.41 | -1.71 | 1.87 | 0.77 |
| Nodakenetin | 26305 | 246.26 | 1 | 1 | 4 | 50.67 | 2.14 | 67.45 | High | 1.15 | 0.38 | -2.82 | 2.04 | 0.64 |
| Phloretin | 4788 | 274.27 | 4 | 4 | 5 | 97.99 | 1.99 | 74.02 | High | -0.32 | -0.92 | -2.53 | 2.38 | 0.21 |
| Isosakuranetin | 160481 | 286.28 | 2 | 2 | 5 | 75.99 | 2.25 | 76.04 | High | 1.1 | 1.9 | -2.17 | 1.84 | 0.11 |
| Pinocembrin | 68071 | 256.25 | 1 | 2 | 4 | 66.76 | 2.26 | 69.55 | High | 1.15 | 0.42 | -2.04 | 1.58 | 0.12 |
| Eriodictyol | 440735 | 288.25 | 1 | 4 | 6 | 107.22 | 1.45 | 73.59 | High | -0.09 | -0.82 | -3.14 | 2.03 | -0.01 |
| Apigenin | 5280443 | 270.24 | 1 | 3 | 5 | 90.9 | 2.11 | 73.99 | High | 1 | -0.73 | -2.06 | 2.45 | 0.56 |
| Chrysin | 5281607 | 254.24 | 1 | 2 | 4 | 70.67 | 2.55 | 71.97 | High | 0.94 | 0.04 | -1.91 | 2.28 | 0.4 |
| Luteolin | 5280445 | 286.24 | 1 | 4 | 6 | 111.13 | 1.73 | 76.01 | High | 0.09 | -0.9 | -2.25 | 2.45 | 0.49 |
| Baicalein | 5281605 | 270.24 | 1 | 3 | 5 | 90.9 | 2.24 | 73.99 | High | 1.11 | -1.06 | -2.21 | 2.32 | 0.25 |
| Oroxylin A | 5320315 | 284.26 | 2 | 2 | 5 | 79.9 | 2.56 | 78.46 | High | 1.02 | -0.11 | -2.21 | 2.41 | 0.31 |
| Daidzein | 5281708 | 254.24 | 1 | 2 | 4 | 70.67 | 2.24 | 71.97 | High | 0.9 | -0.06 | -1.99 | 2.16 | 0.16 |
| Genistein | 5280961 | 270.24 | 1 | 3 | 5 | 90.9 | 2.09 | 73.99 | High | 0.9 | -0.71 | -2.04 | 2.26 | 0.15 |
| Vanillic acid | 8468 | 168.15 | 2 | 2 | 4 | 66.76 | 1.08 | 41.92 | High | 0.33 | -0.38 | -2.62 | 2.45 | 0.62 |
| Caffeic acid | 689043 | 180.16 | 2 | 3 | 4 | 77.76 | 0.93 | 47.16 | High | 0.63 | -0.64 | -2.6 | 2.38 | 0.5 |
| Ferulic acid | 445858 | 194.18 | 3 | 2 | 4 | 66.76 | 1.36 | 51.36 | High | 0.17 | -0.23 | -2.61 | 2.28 | 0.62 |
| Emodin | 3220 | 270.24 | 0 | 5 | 3 | 94.83 | 1.87 | 70.78 | High | 0.05 | -0.72 | -2.33 | 2.11 | 0.34 |
| Shikonin | 479503 | 288.29 | 3 | 3 | 5 | 94.53 | 2.08 | 77.82 | High | 0.59 | -0.6 | -2.46 | 1.59 | 0.07 |
| Ellagic acid | 5281855 | 302.19 | 0 | 4 | 8 | 141.34 | 1 | 75.31 | High | 0.33 | -1.27 | -3.53 | 2.39 | 0.53 |

**Table S5.** Water solubility, Oral rat chronic toxicity (LOAEL), fraction unbound (F_u_) and steady state volume of distribution (VDss) evaluation of the successfully filtered out potential drug molecules.

| Compound name | Ali  logS | ESOL logS | SILICOS-IT  LOGs | Water solubility  (log mol/L) | Oral rat chronic toxicity  (LOAEL) | VDss | F_u_ |
| --- | --- | --- | --- | --- | --- | --- | --- |
| Indole-3-acetamide | -1.58 | -1.78 | -3.27 | -2.12 | 1.4 | 0.15 | 0.39 |
| Antractylenolide III | -2.71 | -2.7 | -3.15 | -3.639 | 1.915 | 0.292 | 0.315 |
| Epiestriol | -3.37 | -3.38 | -2.94 | -3.313 | 1.918 | 0.081 | 0.16 |
| Aloe-emodin | -3.43 | -3.04 | -3.92 | -3.104 | 1.878 | 0.671 | 0.226 |
| Anthrarufin | -4.98 | -4.17 | -4.1 | -2.918 | 2.185 | 0.244 | 0.2 |
| Alizarin | -4.4 | -3.81 | -4.1 | -2.908 | 2.083 | 0.366 | 0.211 |
| Dantron | -4.4 | -3.81 | -4.1 | -2.918 | 2.185 | 0.244 | 0.2 |
| Oxyresveratrol | -4.12 | -3.46 | -2.71 | -3.293 | 1.308 | -0.024 | 0.278 |
| Psoralen | -2.19 | -2.73 | -4.5 | -2.47 | 1.07 | -0.13 | 0.3 |
| Nodakenetin | -2.79 | -2.92 | -3.93 | -3.13 | 1.49 | 0.5 | 0.36 |
| Phloretin | -4.34 | -3.38 | -3.37 | -3.07 | 3.31 | 0.76 | 0.25 |
| Isosakuranetin | -4.1 | -3.7 | -4.12 | -3.09 | 2.14 | 0.21 | 0.07 |
| Pinocembrin | -3.99 | -3.64 | -4 | -3.53 | 2.05 | -0.38 | 0.02 |
| Eriodictyol | -3.9 | -3.26 | -2.84 | -3.25 | 2.47 | 0.37 | 0.1 |
| Apigenin | -4.59 | -3.94 | -4.4 | -3.32 | 2.29 | 0.82 | -2.06 |
| Chrysin | -4.69 | -4.19 | -4.98 | -3.53 | 0.95 | 0.4 | 0.13 |
| Luteolin | -4.51 | -3.71 | -3.82 | -3.09 | 2.4 | 1.15 | 0.16 |
| Baicalein | -4.74 | -4.03 | -4.4 | -3.3 | 2.64 | 0 | 0.15 |
| Oroxylin A | -4.85 | -4.23 | -5.1 | -3.43 | 0.72 | 0.17 | 0.08 |
| Daidzein | -3.53 | -3.6 | -4.98 | -3.79 | 1.18 | -0.17 | 0.1 |
| Genistein | -3.72 | -4.23 | -4.4 | -3.54 | 2.18 | 0.09 | 0.08 |
| Vanillic acid | -2.02 | -2.44 | -1.32 | -1.83 | 2.03 | -1.73 | 0.51 |
| Caffeic acid | -1.86 | -2.38 | -0.71 | -2.33 | 2.09 | -1.09 | 0.52 |
| Ferulic acid | -2.11 | -2.52 | -1.42 | -2.81 | 2.06 | -1.36 | 0.34 |
| Emodin | -3.67 | -4.37 | -3.91 | -3.19 | 2.07 | 0.45 | 0.18 |
| Shikonin | -3.51 | -4.61 | -2.62 | -2.72 | 2.48 | 0.61 | 0.36 |
| Ellagic acid | -2.94 | -3.66 | -3.35 | -3.18 | 2.69 | 0.37 | 0.08 |
